# NaP-TRAP, a novel massively parallel reporter assay to quantify translation control

**DOI:** 10.1101/2023.11.09.566434

**Authors:** Ethan C. Strayer, Srikar Krishna, Haejeong Lee, Charles Vejnar, Jean-Denis Beaudoin, Antonio J. Giraldez

**Author notes:** Corresponding authors (JDB); (AJG, Lead contact).

## Abstract

The *cis*-regulatory elements encoded in a mRNA determine its stability and translational output. While there has been a considerable effort to understand the factors driving mRNA stability, the regulatory frameworks governing translational control remain elusive. We have developed a novel massively parallel reporter assay (MPRA) to measure mRNA translation, Nascent Peptide Translating Ribosome Affinity Purification (NaP-TRAP). NaP-TRAP measures translation in a frame specific manner through the immunocapture of epitope tagged nascent peptides of reporter mRNAs. In contrast to existing MPRA methods, NaP-TRAP does not require specialized equipment and is readily adaptable to steady-state and dynamic model systems. We have employed NaP-TRAP to quantify Kozak strength and the regulatory landscapes of 5’ UTRs in the developing zebrafish embryo and in human cells, characterizing general and developmentally dynamic *cis*-regulatory elements. To this end, we identify U-rich motifs as general enhancers, and upstream ORFs and GC-rich motifs as global repressors of translation. We also observe a translational switch during the maternal-to-zygotic transition, where C-rich motifs shift from repressors to prominent activators of translation. Conversely, we show that microRNA sites in the 5’ UTR repress translation following the zygotic expression of miR-430. Together these results demonstrate that NaP-TRAP is a versatile, accessible, and powerful method to decode the regulatory functions of UTRs across different systems.

---

**L**ife requires spatial and temporal control of protein expression. The protein output of a given transcript reflects the integration of translation and mRNA stability. While there has been a considerable effort to understand the factors driving mRNA synthesis,^1^ maturation,^2^ and decay,^3-6^ the regulatory frameworks governing translational control remain elusive.^7,8^ *Cis*-regulatory elements encoded in the sequence and structure of mRNAs modulate these processes. Although these elements are distributed throughout the transcript, they are often concentrated in the regions up- and downstream of the main coding sequence, the 5’ and 3’ untranslated regions (UTRs) respectively.^9,10^ Given that initiation is the rate limiting step of translation, there has been a particular interest in understanding the regulatory elements found in 5’ UTRs.^11-14^ These elements include internal ribosome entry sites (IRESs),^15^ G-quadruplexes,^16^ iron response elements (IREs),^17^ and upstream Open Reading Frames (uORFs).^18,19^

To investigate translation control, several studies have utilized ribosome profiling.^20,21^ While this method accurately quantifies the translation efficiency of individual genes, its capacity to characterize the function of *cis*-regulatory elements is limited. The generation of ribosome protected fragments decouples the translation measurement of a given mRNA from its cognate untranslated regions.^22^ Thus, measurements of translation efficiency often reflect the amalgamation of several isoforms of a given gene, each with a unique set of regulatory elements. This process is complicated further by the fact that endogenous transcripts contain multiple regulatory elements that exert differential and often competing effects on translation.

Massively parallel reporter assays (MPRAs) are particularly suited to address these challenges. MPRAs measure the abundance and/or translation efficiency of thousands of reporters simultaneously. These assays have improved our understanding of translation control and mRNA stability. Previous works have characterized Kozak strength,^23^ uORFs,^24-26^ IRESs,^27,28^ codon optimality,^29,30^ RNA structure,^31^ microRNA binding sites,^32^ and the effects of variation in human UTRs.^33-36^ These assays have also identified novel sequence motifs driving translation and decay.^4,37,38^ Despite these insights, conclusions based on MPRAs have been limited by the methods used to measure translation: growth-selection,^39^ fluorescent-cell sorting (FACS),^24,40^ polysome profiling,^34,36^ and direct analysis of ribosome targeting (DART)^38^ (Table 1). In MPRAs where reporters are encoded in DNA, measurements of translation can be confounded by additional layers of transcriptional regulation. In polysome fractionation, which quantifies the number of ribosomes on each transcript, inactive ribosomes, ribosomes translating out of frame, or ribosomes translating ORFs outside of the coding sequence may skew translation measurements. Additionally, the methodological complexity of these assays has reduced the use of translation-based MPRAs across diverse systems.

**Table 1.**
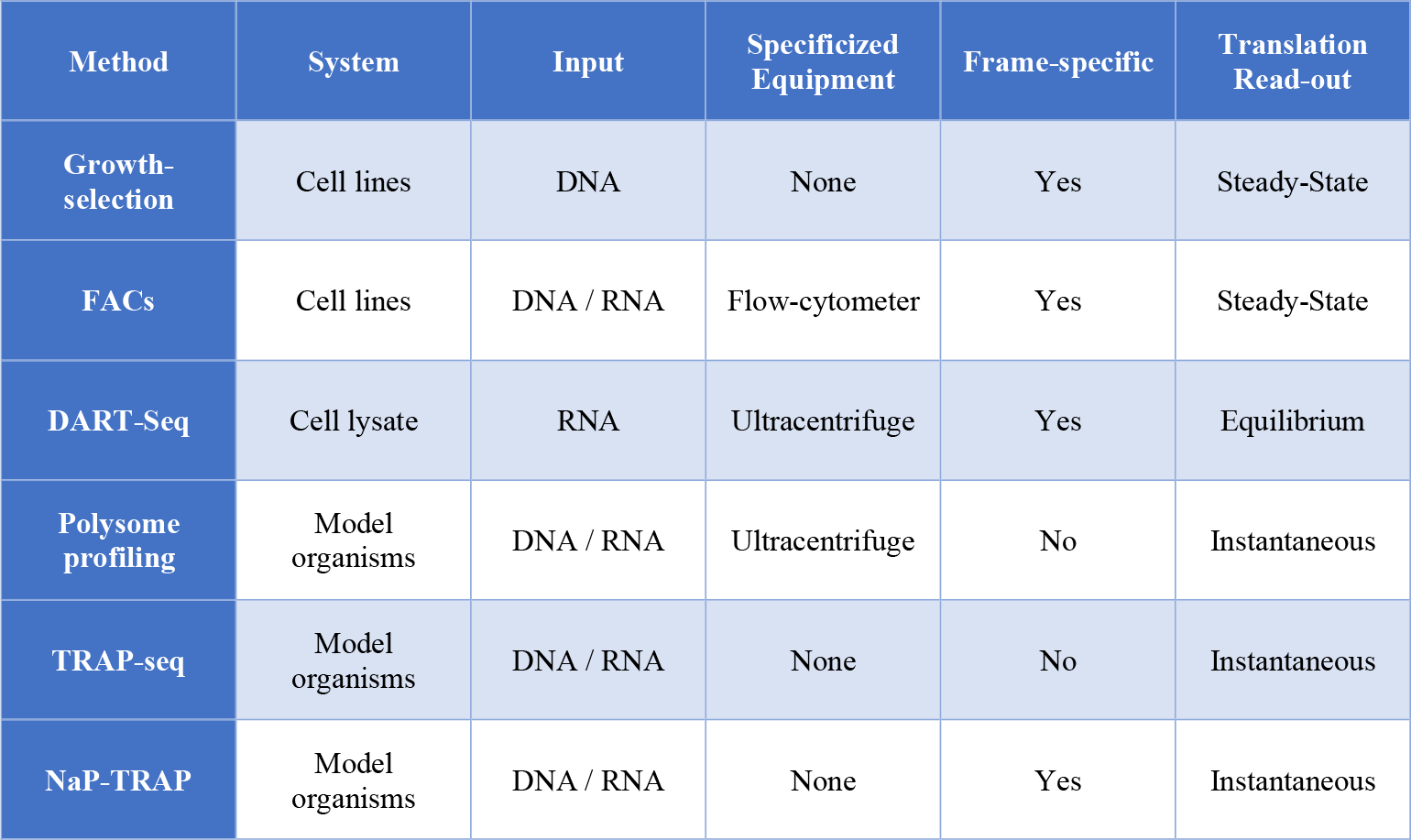
Comparing NaP-TRAP to existing translation based MPRA methods. NaP-TRAP was developed to quantify the translation of thousands of reporters simultaneously in a frame-specific manner. In contrast to existing methods, NaP-TRAP can be adapted to dynamic model systems (e.g. the developing zebrafish embryo), and does not require specialized equipment or a large amount of source material to measure translation.

Here, we develop NaP-TRAP (Nascent Peptide Translating Ribosome Affinity Purification), a novel method to measure in-frame translation through the immunocapture of nascent peptides. More specifically, by including an N-terminal FLAG tag in the coding sequences of reporter mRNAs, we enrich for reporters in a manner proportional to the number of active ribosomes translating the main ORF in frame. We validate NaP-TRAP using reporter assays of increasing complexity. First, we measure the translation of individual reporters containing known *cis*-regulatory elements. Next, we quantify Kozak strength in zebrafish by measuring the translation of thousands of reporters simultaneously. Lastly, we assess the regulatory potential of endogenous 5’ UTR sequences in the developing zebrafish embryo and HEK293T cells to identify common and developmentally modulated motifs that regulate translation in vertebrates. In doing so we demonstrate that NaP-TRAP is an accessible, versatile, and quantitative method, which has the capacity to measure translation control across multiple systems.

## Results

### NaP-TRAP measures translation via the immunocapture of nascent chains

*Cis*-regulatory elements encoded in mRNAs modulate translation efficiency. To measure the regulatory potential of these elements we developed a novel massively parallel reporter assay (MPRA), NaP-TRAP (Nascent Peptide Translating Ribosome Affinity Purification). We reasoned that we could enrich for reporters in a manner proportional to their translation efficiency through the immunocapture of FLAG-tagged nascent chain complexes immobilized by cycloheximide treatment (Figure 1A). To this end, we measured translation as a ratio of the amount of a reporter mRNA in the pulldown relative to its input. This approach decouples translation measurements from differences in mRNA abundance.

**Figure 1.**
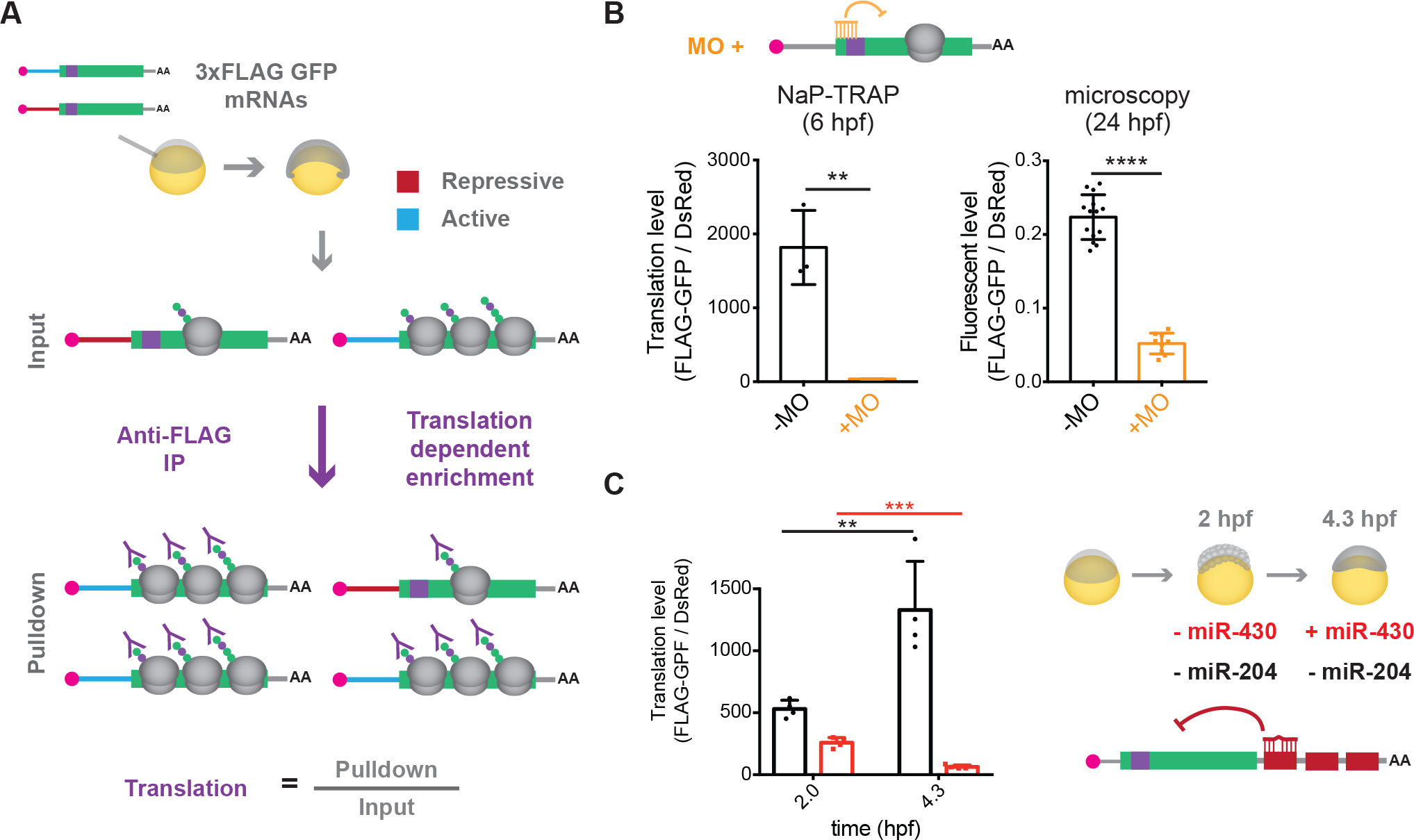
NaP-TRAP measures translation through immunocapture. (A) Schematic detailing the NaP-TRAP method. FLAG-tagged nascent chain complexes of reporter mRNAs are enriched via an anti-FLAG immunoprecipitation. Translation is measured as a ratio of reporter reads in the pulldown relative to the input. (B) NaP-TRAP derived translation at 6 hpf (left, N = 3) and immunofluorescence measurements at 24 hpf (right, -MO N = 14, +MO N = 9) in the presence or absence of a translation blocking morpholino (unpaired t-test: ** p < 0.005, **** p < 0.0001). (C) NaP-TRAP derived translation values for 3xFLAG-GFP-3xmiR-430 (red) and 3xFLAG-GFP-3xmiR-204 (black) at 2 hpf (N = 4) and 4.3 hpf (N = 4) (unpaired t-test: ** p < 0.01, *** p < 0.005). Schematic representation of repression of translation exercised by miR-430, but not miR-204 at 4.3 hpf.

To evaluate the capacity of NaP-TRAP to measure translation we performed two reporter assays. First, we measured the effect of a translation blocking morpholino. We co-injected 3xFLAG-GFP and dsRED mRNAs into single cell zebrafish embryos in the presence or absence of a translation blocking morpholino, targeting the start codon of 3xFLAG-GFP and performed NaP-TRAP at 6 hpf. To measure the amount of 3xFLAG-GFP reporter mRNA present in the input and pulldown fractions of each anti-FLAG immunoprecipitation, we employed reverse transcription-quantitative polymerase chain reaction (RT-qPCR). Given that the dsRED reporter mRNA was neither enriched by the anti-FLAG immunoprecipitation nor targeted by the translation blocking morpholino, we utilized the relative abundance of the dsRED reporter mRNA in each fraction to normalize NaP-TRAP derived translation values across experimental conditions. Using NaP-TRAP we observed a 48-fold decrease in translation in the presence of the morpholino at 6 hours post fertilization (hpf) (Figure 1B). These results are consistent with the translational repression in morpholino injected embryos observed by the decrease in fluorescence (GFP/dsRED) at 24 hpf (Figure 1B). Next, we tested the capacity of NaP-TRAP to capture dynamic translation control. In the developing zebrafish embryo, the microRNA miR-430 is one of the first zygotically transcribed genes. At 4.3 hpf (hours post fertilization), the expression of miR-430 results in the translation repression, deadenylation, and eventual decay of targeted mRNAs.^41,42^ To quantify this effect, we measured the translation of 3xFLAG-GFP reporters with partial complementarity to two different microRNAs in their 3’ UTRs, miR-430 (3xFLAG-GFP-3xmiR-430) and miR-204 (3xFLAG-GFP-3xmiR-204). We selected miR-204 target sites as a control given that this microRNA is not expressed in the early embryo. Using NaP-TRAP we measured the translation of mRNA reporters at 2 and 4.3 hpf, before and after miR-430 expression. As described above we utilize the abundance of the dsRED control RNA to normalize translation values across experimental conditions. Between the two timepoints, the translation of the 3xmiR-430 reporter decreased by ∼4.3 fold, whereas the translation of the 3xmiR-204 reporter increased by ∼2.5 fold (Figure 1C). These results are consistent with the role of miR-430 in translational repression and the global increase in translation observed during the early stages of development.^43^ Together these results demonstrate that NaP-TRAP measures the translation of individual reporters targeting general and developmentally dynamic *cis*-regulatory elements. Further, these experiments highlight the versatility of the method as they demonstrate the capacity of NaP-TRAP to quantify *cis*-regulation mediated by the 5’ and 3’ UTRs.

### Using NaP-TRAP to investigate Kozak strength in the developing zebrafish embryo

Next, to evaluate the capacity of NaP-TRAP to function as a MPRA, we designed a library to quantify the regula-tory potential of the Kozak sequence. We elected to measure Kozak strength for two reasons: (1) the Kozak sequence is a strong determinant of translation initiation and (2) Kozak strength is often inferred based on the frequency of these sequences in the genome.^44,45^ To generate a library of diverse Kozak sequences, we incorporated six random nucleotides upstream and one random nucleotide downstream of the AUG of the 3xFLAG-GFP reporter (Figure 2A).^46^ We injected the *in vitro* transcribed mRNA library into single cell zebrafish embryos and measured the translation using NaP-TRAP at 6 hpf. We observed a strong correlation in translation values between replicates (Pearson’s r = 0.78, Figure S1A). To identify active and repressive Kozak sequences, we generated position weight matrices for the top and bottom 10 % of reporters based on their translation (Figure 2A). We observed that active Kozak sequences were enriched in C’s and A’s, whereas repressive Kozak sequences were enriched in U’s and G’s.

**Figure 2.**
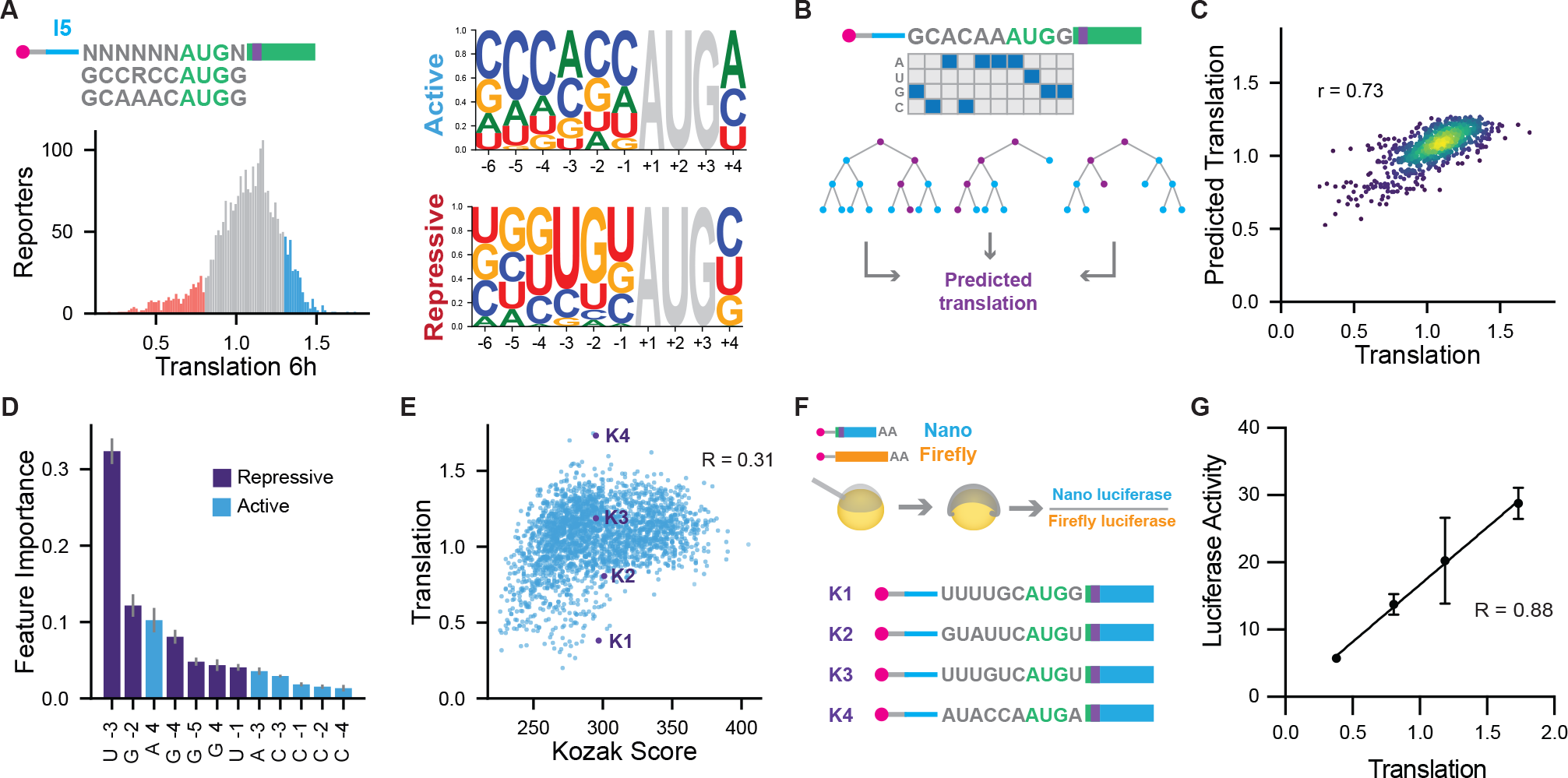
NaP-TRAP measures the effect of Kozak strength on translation. (A)A schematic detailing the Kozak library containing six random nucleotides upstream and one nucleotide downstream of the start codon of 3xFLAG-GFP (top left). Histogram of NaP-TRAP translation values at 6 hpf (bottom left). Sequence logos for the top and bottom 10% of reporters based on translation (right).^51^ (B,C) Cartoon of random forest regression model feature generation through the one-hot-encoding of reporter sequences (B). Scatterplot comparing the model’s prediction to the experimentally derived translation values of a test set of reporters (C; Pearson’s R) (N = 785 reporters). (D) Permuted feature importance derived from random forest regression model. Nucleotide positions in purple correlate negatively with translation, whereas positions in blue correlate positively with translation (Spearman Rank Correlation Coefficient). (E) Comparison of translation measurements at 6 hpf and an in silico derived Kozak score based on the frequency of Kozak sequences in the transcriptome (Pearson’s R) (N = 2,617 reporters).^23^ (F,G) The translation of four reporters with a Kozak score between 295-305 was measured using a dual luciferase-based assay (F). Plot comparing relative luciferase activity (Nano luciferase / Firefly luciferase), and NaP-TRAP translation values (G) (Pearson’s R, N = 3).

To examine the effect of nucleotide identity and position on translation we employed a random forest regression model (RFM). Model features were generated based on nucleotide identities and positions within the Kozak sequence (Figure 2B). Prior to training the model, we divided the data into a test and a training set, 30% and 70% of reporters respectively. Using the training set, we employed a 5-fold cross validation to optimize model parameters. The model’s prediction explained 73% of the variance observed in the test set data (Figure 2C). Given this strong correlation we employed the model to predict the translation of all possible Kozak sequences (Figure S1C, Table S2). To identify the features informing the predictive power of the model, we performed a permutated feature importance analysis. Briefly, we shuffled the identities of the features of the model one at a time and measured the effect of the change on the predictive power of the model. At 6 hpf bases G, G, U, and G in positions -5, -4, -3, and -2 were the most repressive features, whereas C, A/C, and A in positions -4, -3, +4 were the most activating features (Figure 2D and S1B).

Lastly, we compared NaP-TRAP translation values to an i*n silico* derived metric, Kozak score. In zebrafish Kozak strength has been previously inferred based on the frequency of a given Kozak sequence within the transcriptome. This metric associates Kozak sequence abundance with increased translation initiation.^45^ Our experimentally derived translation values challenge this hypothesis as we observed a weak correlation between NaP-TRAP and Kozak score (R = 0.31, Figure 2E). To validate this conclusion, we identified sequences that had similar Kozak scores (300 +/-5) yet exhibited different NaP-TRAP derived translation values and measured their translation using a dual luciferase reporter assay (Figure 2F). The luciferase translation values correlated strongly with NaP-TRAP (Pearson’s R 0.88; Figure 2G). This result not only highlights the importance of measuring Kozak strength experimentally, but also demonstrates that NaP-TRAP derived translation values correlate well with protein production. Taken together, these results demonstrate that NaP-TRAP can be employed as a MPRA. By measuring the translation of thousands of reporters simultaneously, we quantify and model Kozak strength in a vertebrate system.

## Investigating the regulatory potential of endogenous 5’ UTRs

Upon validating NaP-TRAP’s capacity to function as a MPRA, we investigated the regulatory complexity of endogenous 5’ UTRs during the early stages of embryogenesis. Prior to zygotic genome activation, the translation of maternally supplied mRNAs drives development.^47,48^ Given that initiation is the rate limiting step of translation, 5’ UTRs provide a mechanism for maternally supplied mRNAs to modulate their protein output. To study the *cis*-regulatory elements encoded in these mRNAs, we designed an 11,088-sequence synthetic oligo library (124-nt long; Table S3) by tiling the 5’-UTRs of 1,725 zebrafish genes every 25 nu-cleotides (Figure 3A). As we were interested in identifying specific sequence elements, which modulated differential translation regulation we elected to keep the 5’ UTR length and Kozak sequence of reporter mRNAs constant as both of these features correlate strongly with translation efficiency in the developing embryo. We injected the *in vitro* transcribed library into single cell zebrafish embryos and measured translation at 2 hpf and 6 hpf using NaP-TRAP. The addition of an internal spike-in at the RNA extraction step allowed us to quantify relative changes in translation across developmental stages (Figure 3A). Translation values were strongly correlated across replicates (Pearson’s R >= 0.90; Figure S2A-B, S3A-B), and with protein abundance at both time points (dual luciferase assay; 2 hpf Pearson’s R 0.75, 6 hpf Pearson’s R 0.88 Figure S2C-D). We observed a mean increase in translation between 2 hpf and 6 hpf of ∼1.9 fold, consistent with the gradual de-repression of translation during the first 24 hours of development (Figure 3B).^43,49^

**Figure 3.**
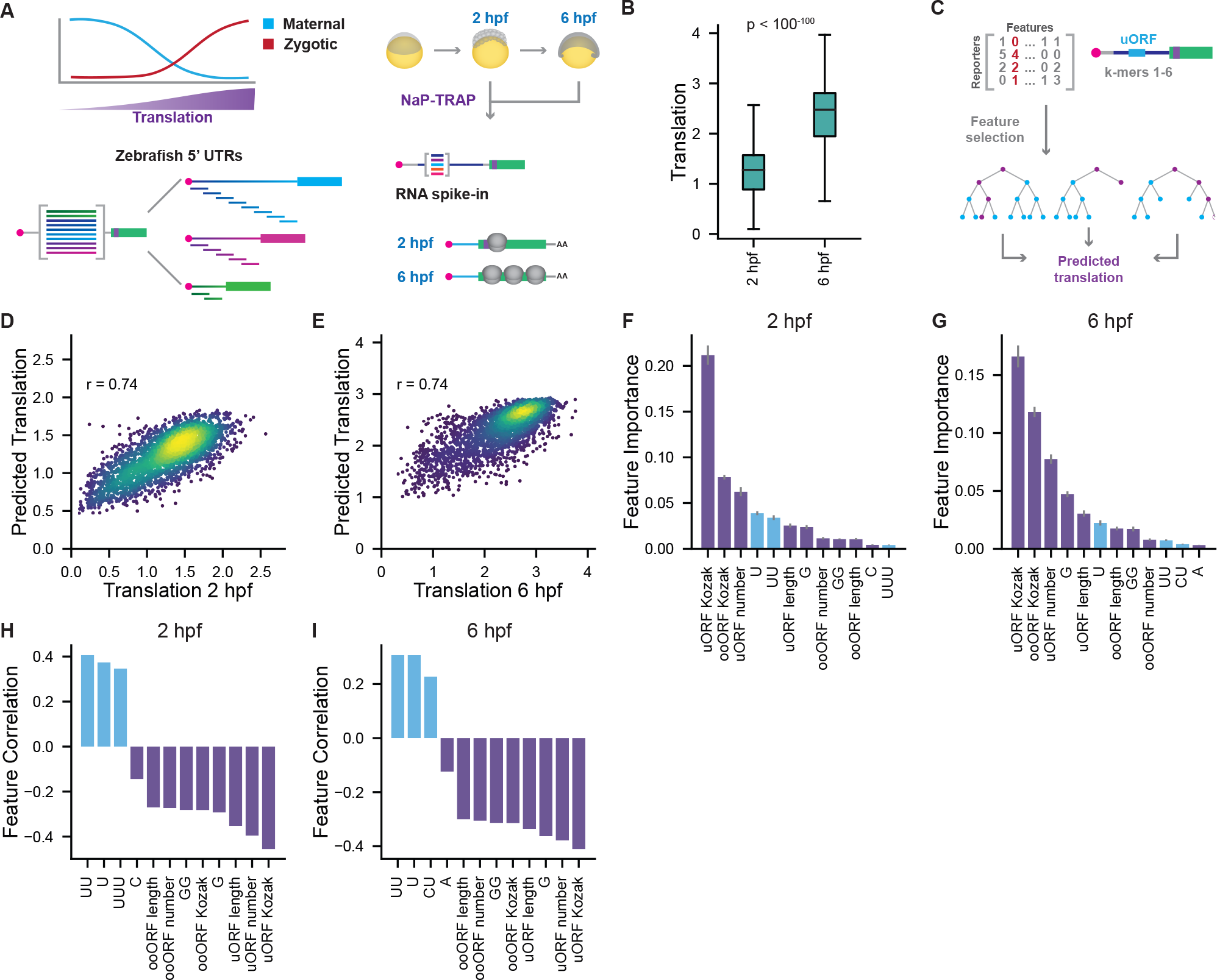
Using NaP-TRAP to predict the translation of zebrafish 5’ UTRs. (A) A schematic detailing global changes in translation during the maternal-to-zygotic transition in the developing zebrafish embryo (top left). The 5’ UTR library was generated by tiling the 5’ UTRs of zebrafish (bottom left). The NaP-TRAP workflow with the addition of spike-ins at the RNA extraction step (right). (B) Comparison of translation values between 2 and 6 hpf of the 5’ UTR library (Mann-Whitney p < 10^-100^, N = 8,529 reporters). (C) Schematic detailing random forest regression model feature selection (k-mer counts 1-6 nt and features characterizing uAUGs). (D-I) Scatterplot comparing each model’s prediction to the experimentally derived translation values of a test set of reporters (N = 2,559 reporters) (D,E; Pearson’s R). The permuted feature importance of features (top 12) with the greatest effect on model performance (blue refers to a positive correlation, purple refers to a negative correlation; Spearman Rank Correlation Coefficient) (F,G). The correlation of selected features with translation at the indicated timepoints (H,I).

Next, we utilized a random forest regression model trained on sequence elements and features characterizing upstream AUGs (uAUGs) to predict translation (Figure 3C and S3D-I). These models explained 74% of the variance observed in the test set data at both timepoints (Figure 3D and 3E). Using a permuted feature importance analysis, we identified the number of upstream uORFs (uORFs) as well as the Kozak strength of uORFs and out of frame overlapping ORFs (ooORFs) as the features contributing most significantly to the predictive power of the model at both timepoints (Figure 3F and 3G). When we examined the correlation of selected features with translation, we observed that U-repeats activated translation and G-repeats and uORFs suppressed translation at both timepoints. Interestingly, we also observed that C-rich k-mers suppressed translation at 2 hpf, whereas at 6 hpf C- and CU-rich k-mers activated translation (Figure 3H and 3I). Given the prominent role of upstream open reading frames in the feature importance analysis, we repeated the random forest analysis on reporters that contained no upstream start codons (Figure S2E-J). In the absence of upstream start codons, the importance and prominence of C- and CU-rich k-mers increase at 6 hpf (Figure S2J). Altogether these results demonstrate the following: (1) uAUGs are the most prominent repressive elements during early embryogenesis, and (2) 5’ UTR regulatory landscapes are dynamic during development.

### Identifying sequences driving differential translation

To identify the sequence elements modulating differential translation during development we divided the 5’ UTR reporters into four groups based on their relative translation at 2 and 6 hpf: (1) repressed, (2) active, (3) repressed post ZGA, and (4) active post ZGA (Figure 4A and S3J). Next, we performed a differential pentamer enrichment analysis on each group relative to the reporter library. This analysis revealed that repressed reporters were significantly enriched in upstream ORFs and GC-rich pentamers and depleted in U-rich tracks (p >10^-5^ hypergeometric test, Fig-ure 4B and S3K). Conversely, active reporters were enriched in U-rich pentamers (UUUUU, CUUUU, UUUUA, GUUUU) and depleted in upstream start codons (Figure 4C and S3L). Reporters that were more highly translated after genome activation were enriched in C-rich pentamers (CU-CUC, CUCCC, CCAUC, CCUCC) and depleted in U-repeats (Figure 4D and S3M). In contrast, reporters that were repressed post ZGA were enriched in AG/UG-rich sequences (UAGUG, UAUUG, AAGAA, AGACU), as well as sequence motifs complementary to the seed site of the microRNA miR-430 (GCACU and GCACUU; Table S3) and depleted in uORFs and pyrimidine repeats (Figure 4E and S3N).

**Figure 4.**
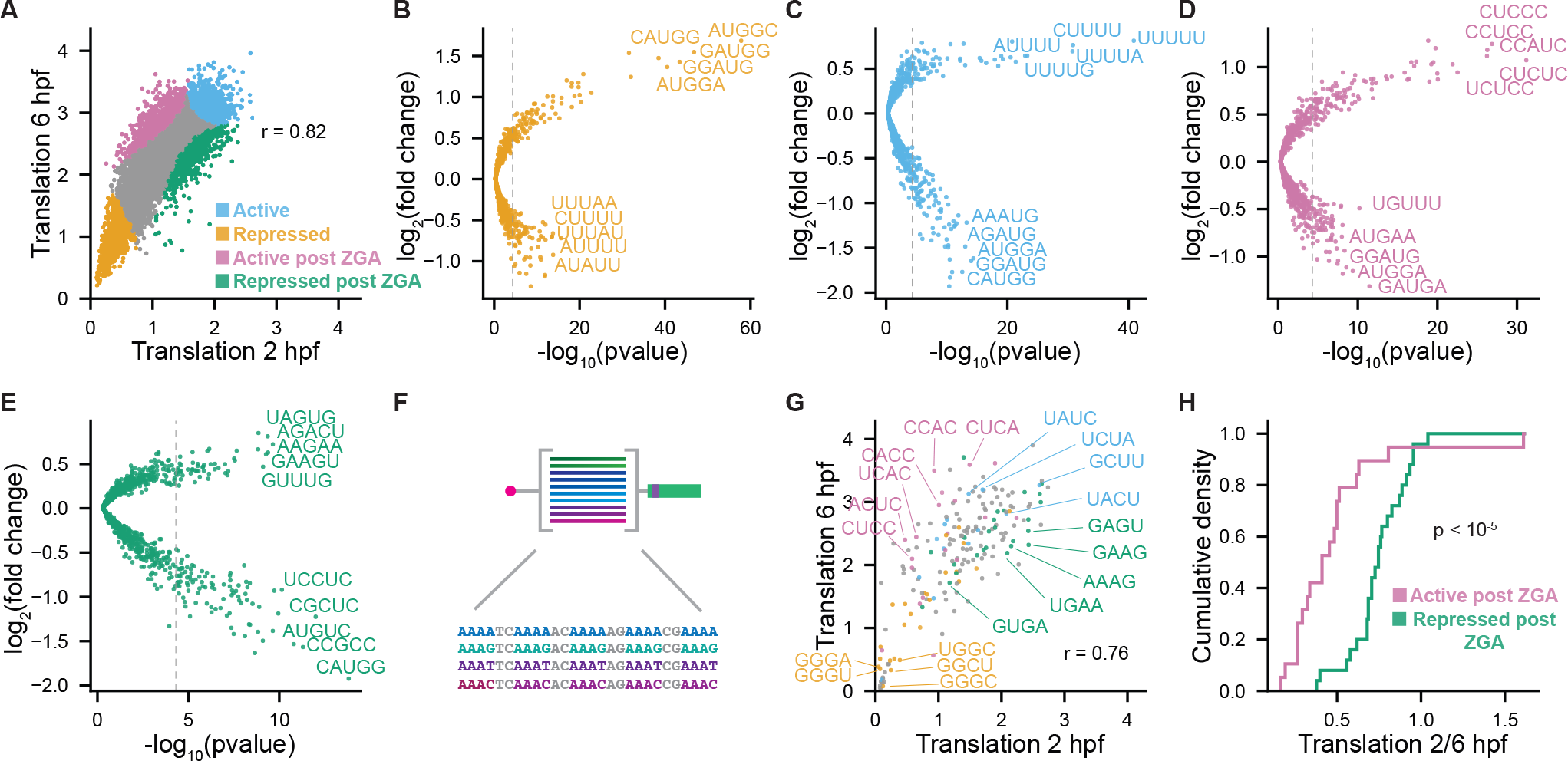
NaP-TRAP captures general and dynamic 5’ UTRs *cis*-regulation. (A)The translation of the 5’ UTR library at 2 hpf and 6 hpf using NaP-TRAP (Pearson’s R, N = 8,529 reporters). Reporters were divided into four groups based on their translation rank at both time-points: repressed, active, active post ZGA, and repressed post ZGA (orange, blue, pink, and green, respectively). (B-E) Fold-change enrichment and depletion for all pentamers in each group from (A) relative to the reporter library (hyper-geometric test to calculate significance; Bonferroni corrected p-value threshold p < 5 * 10^-6^). (F) Schematic detailing the library of all possible tetramer repeats separated by dinucleotide spacers. (G) Translation measurements of the validation library at 2 and 6 hpf (Pearson’s R, N = 195 reporters). Tetrameric reporters were labeled based on whether their repeat was enriched in the reporter groups described above (A). (H) Cumulative distribution plot comparing the translation (2hpf / 6 hpf) of validation reporters identified as active post ZGA (pink) and repressed post ZGA (green) (Mann-Whitney U-test: p < 10^-5^).

To validate these results, we generated a library of 5’ UTR sequences containing all possible tetramer repeats separated by dinucleotide spacers and measured translation using NaP-TRAP at 2 and 6 hpf (Figure 4F). Next, we plotted the translation values of the validation library labeling each reporter on the basis of whether its encoded tetramer repeat was enriched in the reporter groups described above (Figure 4G and Table S3 and Table S4). The distributions of reporters encoding repressed, repressed post ZGA, and active post ZGA tetramers largely recapitulated the distributions of their respective groups in the 5’ UTR library (Figure S4B-E). Consistent with this, a cumulative analysis of the translation (2 hpf / 6 hpf) of reporters enriched in “active motifs post ZGA” revealed a larger increase in translation compared to those enriched in “repressed motifs post ZGA” (Figure 4H). Altogether these results suggest that there are general and dynamic sequence motifs in zebrafish 5’ UTRs that modulate translation during the early stages of development.

### Complementary sequences to miR-430 in the 5’ UTR suppress translation

miR-430 is one of the first zygotically transcribed genes in the developing zebrafish embryo.^41^ The enrichment of miR-430 seeds in reporters that were repressed after ZGA suggests that the maternal-to-zygotic transition is shaped by 5’ UTR mediated translation control. To determine whether this repressive effect is specific for miR-430, we compared the translation between 2 hpf and 6 hpf for reporters containing seeds for miR-430 or miR-1, a microRNA that is not expressed in the early stages of development. We observed a significant decrease in the translation of miR-430 containing reporters (6 vs 2 hpf) relative to the translation of reporters lacking either seed. In contrast, there was a slight increase in translation of miR-1 control reporters (Figure 5A). To investigate the mechanism driving miR-430 mediated translation repression, we measured the degree of complementarity between the microRNA and the targeted reporter. Translation of the miR-430 reporters was negatively correlated to miRNA-5’ UTR complementarity (Figure 5B). Whereas translation of the miR-1 control reporters was not significantly correlated with miRNA-5’ UTR complementarity (Figure 5C). This observation suggests that 5’ UTR microRNA mediated translation repression is driven by sequences with homology beyond the seed site of the microRNA.

**Figure 5.**
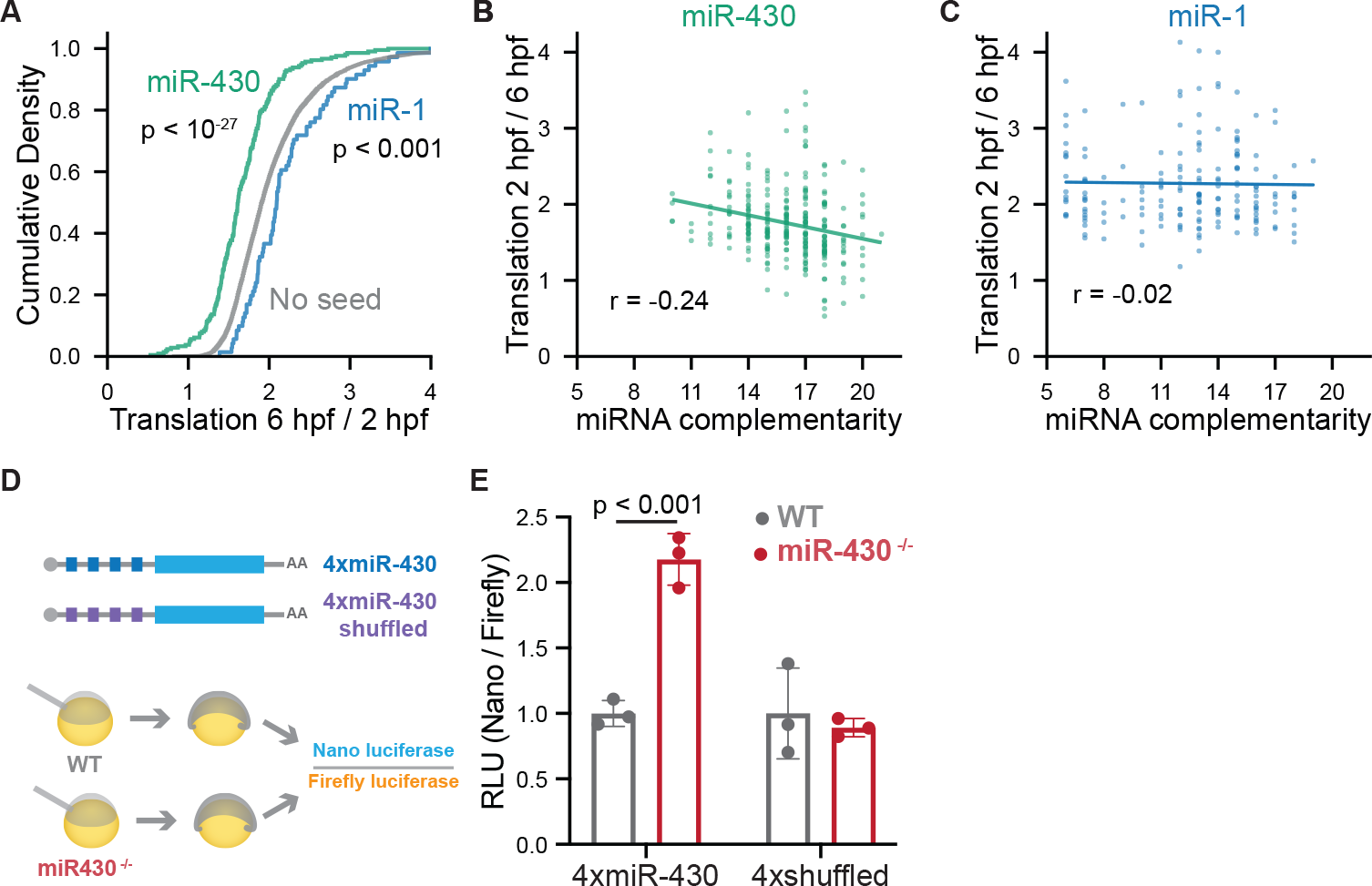
Zygotic expression of miR-430 inhibits translation of mRNAs with 5’ UTR seed sites. (A) Cumulative distributions of translation (6 hpf / 2 hpf) for reporters containing miR-430 (GCACUUA, GCACUUU, AGCACUU; N = 212 reporters) or miR-1 (ACAUUCC, CAUUCCA; N = 71 reporters) heptamers, labeled in green and blue respectively. Reporters labeled in gray contain neither seed sequence (N = 8,242 reporters) (Mann-Whitney U test, p < 10^-27^ miR-430 reporters vs reporters with no seed; Mann-Whitney U test, p < 0.001 miR-1 reporters vs reporters with no seed). (B and C) The effect of number of complementary bases to miR-430 (B, N = 275 reporters) or miR-1 (C, N = 178 reporters) on the translation (6 hpf / 2 hpf) of reporters with seed sequences (Pearson’s R). (D and E) Schematic detailing dual luciferase assay measuring the inhibitory effect of miR-430 binding sites in the 5’ UTR (D). 4xmiR-430-nanoluc and 4xmiR-430-shuffled-nanoluc were injected in wild-type and miR-430 knockout embryos. Relative Luciferase Activity (RLU) values were normalized to each reporter (N = 3, unpaired t-test) (E).

To demonstrate further that miR-430 expression drives the translation repression of 5’ UTRs with seed sites, we constructed two nano-luciferase reporters: 4xmiR-430-nanoluc and 4xmiR-430-MUT-nanoluc. We shuffled the miR-430 sites in the 4xmiR-430-MUT reporter to prevent miR-430 targeting (GCACUU to GCUCUA). We injected these reporter mRNAs together with a firefly luciferase control mRNA into wild-type and *miR-430* ^-/-^ mutant embryos and measured relative luciferase activity at 6 hpf (Figure 5D). We observed a ∼2-fold decrease in the relative luciferase activity of the 4xmiR-430 reporter in the wild-type condition when compared to the mutant. In contrast, we observed no significant difference in luciferase activity between the shuffled reporters when comparing the mutant and wild-type embryos (Figure 5E). This finding suggests that the zygotic expression of microRNA miR-430 represses the translation of mRNAs with 5’ UTR seed sites. Taken together these results demonstrate that miRNAs that target sites in the 5’ UTR can provide significant translational repression *in vivo*.

### The developmentally dynamic role of C-rich motifs

We also observed an enrichment of C-rich pentamers in reporters that were active following ZGA. To explore this observation in greater detail, we measured the mean difference in translation between reporters enriched and depleted in each feature (k-mers ≤ 4) that lacked upstream open reading frames. Next, we compared the rank of features with a significant difference in translation at 2 hpf and 6 hpf. We observed that the k-mers C, CC, CCC, and UCC were some of the most repressive features at 2 hpf yet activated translation at 6 hpf (Figure 6A). To investigate this differential regulation further, we selected two active (U-rich) and two active post ZGA (C-rich) reporters from the 5’ UTR library. To determine the role of C-rich sequences in this translation regulation we mutated U’s in the active reporters to C’s and mutated C’s in the active post ZGA reporters to U’s for a total of eight reporters: four wild-type and four mutants, respectively. We injected the reporters into single cell zebrafish embryos and measured translation at 2 hpf and 6 hpf using NaP-TRAP. At 2 hpf we observed that mutating C’s to U’s in the C-rich reporters enhanced translation (Figure 6C), whereas mutating U’s to C’s in U-rich reporters repressed translation (Figure 6D). In contrast, this effect was largely reduced at 6 hpf (Figure 6E and 6F). These results support the findings of the differential enrichment analysis and indicate that there is a translational switch driven by C-rich sequences following ZGA (Figure 6B). Altogether, these results validate novel developmentally dynamic mechanisms of 5’ UTR mediated translation control, demonstrating the importance of quantifying *cis*-regulation in non-steady state systems.

**Figure 6.**
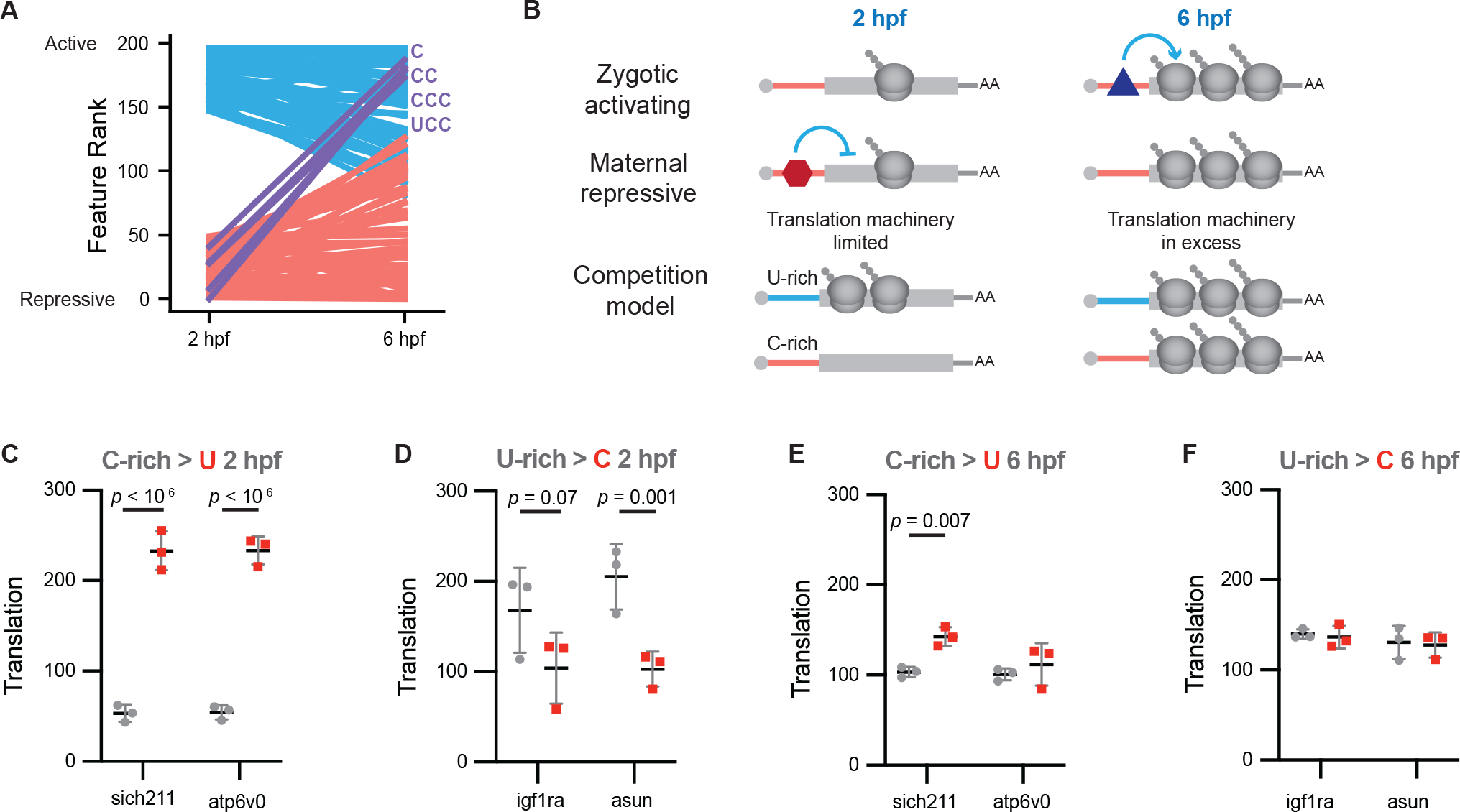
The dynamic effect of C-richness on translation. (A) Plot comparing feature rank at 2 and 6 hpf (k-mers ≤ 4). At each time point feature rank was determined by calculating the mean difference in translation between non-uAUG reporters enriched and depleted (top and bottom 20%) in each feature (red: repressive at 2 hpf, blue: active at 2 hpf, purple: features that were repressive at 2 hpf and then active at 6 hpf). Features were only included if there was a significant difference in mean translation at each timepoint (Bonferroni corrected t-test). (B) Proposed models explaining how the maternal-to-zygotic transition can alter the effect of *cis*-regulatory elements in the 5’ UTR. (C-F) U’s in two active (U-rich) reporters were mutated to C’s (U > C), whereas C’s in two active post ZGA (C-rich) reporters were mutated to U’s (C > U). Translation values were measured using NaP-TRAP at 2 and 6 hpf. Wild-type reporters are colored gray, whereas mutant reporters are labeled red (N = 3, unpaired *t*-test).

### NaP-TRAP can be readily adapted to human cells

To demonstrate the broad applicability of our method, we adapted NaP-TRAP to human cells. To this end, we transfected the *in vitro* transcribed mRNA 5’ UTR library into HEK293T cells and measured translation at 12 hours post transfection (hpt) using NaP-TRAP (Figure 7A and 7B). Replicates were strongly correlated (Pearson’s R ≥ 0.92, Figure S5A-C). To identify motifs modulating translation in HEK293T cells we performed a differential enrichment analysis on active and repressed reporters. Repressed reporters were enriched in AUG-containing motifs and depleted in U-rich motifs (Figure 7C), whereas active reporters were enriched in C-rich pentamers and depleted in AUG containing motifs (Figure 7D), consistent with those observed in zebrafish at 6 hpf (Figure 7E and S5D).

**Figure 7.**
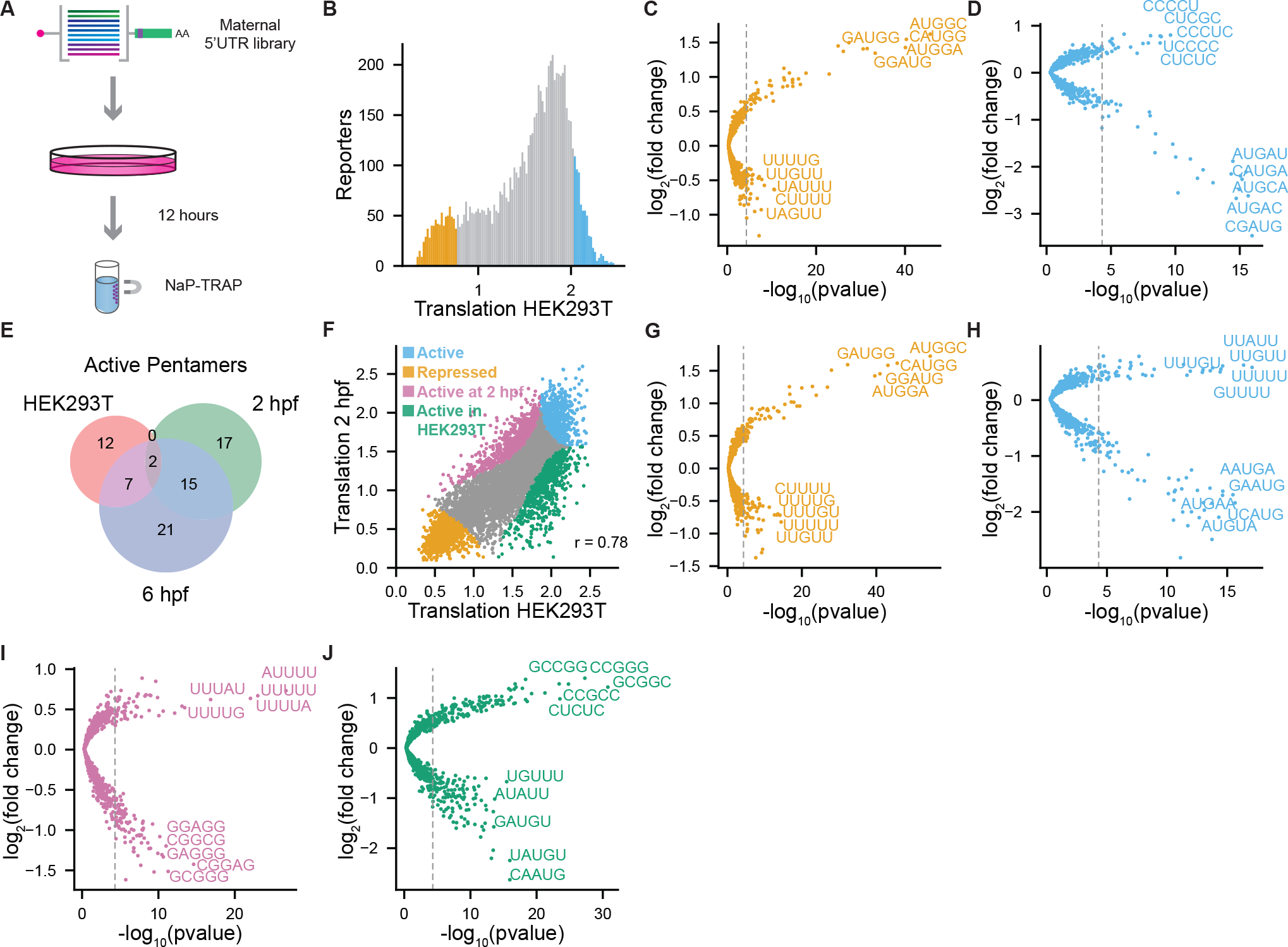
Adapting NaP-TRAP to human cell lines. (A) Schematic detailing NaP-TRAP in HEK293T cells. Cells were transfected with the 5’ UTR library using lipid nanoparticles. Translation was measured at 12 hours post transfection. (B) The distribution of translation values of the 5’ UTR library in HEK293T cells. The top and bottom 10% of reporters based on their translation values are labeled in orange and blue (repressed and active reporters, respectively; N = 7,506 reporters). (C and D) A differential enrichment analysis identified pentamers enriched and depleted in repressed (C) and active (D) reporters, blue and orange respectively (hypergeometric test with a Bonferroni corrected p-value). (E) Venn-diagram comparing the pentamers enriched in active reporters in HEK293T cells and zebrafish embryos at 2 and 6 hpf. (F) Translation values at 2 hpf in zebrafish compared to translation values in HEK293T cells (Pearson’s R). Reporters were divided into four groups based on their translation rank at each condition: active (blue), repressed (orange), active in zebrafish at 2 hpf (pink) and active in HEK293T cells (green) (N = 7,506 reporters). (G-J) The fold-change enrichment or depletion for all pentamers in each group (active (G), repressed (H), active in zebrafish at 2 hpf (I), and active in HEK293T (J)) relative to the reporter library. Significance was calculated using a hyper-geometric test (Bonferroni corrected p-value threshold).

Next, we performed a differential enrichment analysis comparing the human data to each of the zebrafish timepoints (Figure 7F-J and Figure S5E-I, 2 hpf and 6 hpf, respectively). We divided the reporters into four groups: (1) repressed, (2) active, (3) active in zebrafish, and (4) active in HEK293T cells. From these analyses, we conclude that uAUGs are general repressors of translation (Figure 7G and S5F) and U-rich motifs are general activators of translation (Figure 7H, S5G and S5L). We also observed that reporters that are active only in HEK293T cells are enriched in G-rich k-mers, whereas reporters active at 2 hpf or 6 hpf are depleted in these motifs (Figure 7J and S5I). Next, when we employed a random forest model to predict translation. The model explained 80% of the variation in the experimental data (Figure S5J). Consistent with the results observed in zebrafish, Kozak strength and number of upstream AUGs were the most predictive features of translation in HEK293T cells, exhibiting a strong negative correlation with translation (Figure S5K and S5L). All together these results demonstrate that (1) NaP-TRAP is a robust method that can measure translation across multiple model systems, (2) upstream AUGs are a dominant driver of 5’ UTR mediated translation control in HEK293T cells, and (3) U-rich motifs are general activators of translation.

## Discussion

Here, we develop a novel MPRA method, NaP-TRAP, and demonstrate its capacity to measure translation quantitatively. We use NaP-TRAP to characterize *cis*-regulatory elements in the 5’ and 3’ UTRs, the strength of the Kozak sequence, and the regulatory landscape of the 5’ UTRs of mRNAs in the developing zebrafish embryo and HEK293T cells. Using NaP-TRAP we have identified general and developmentally dynamic *cis*-regulatory elements, as well as characterized global changes to translation associated with early embryogenesis.

NaP-TRAP is an accessible, versatile, and quantitative MPRA method to measure translation. In contrast to existing approaches NaP-TRAP does not require a large amount of input material or specialized equipment, making the method adaptable to a wide range of model systems, different cell types, and physiological states (Table 1). Further, NaP-TRAP measures frame-specific translation as only the nascent chains of ribosomes translating the main ORF are FLAG-tagged and thereby immunocaptured. This approach is particularly important in systems with a low basal level of translation (e.g., the early stages of vertebrate embryogenesis^43^ and neurons^50^) and results in translation measurements that reflect protein output (Figure 2G and S2C-D). In polysome profiling and other TRAP methods, inactive ribosomes or ribosomes translating outside of the main open reading frame may affect the measurements of translation. By enriching for reporters through the immunocapture of FLAG-tagged nascent chain complexes, NaP-TRAP measures translation in a manner that is proportional to the number of ribosomes actively translating the tagged ORF. Second, by injecting or transfecting *in vitro* transcribed mRNA reporter libraries, NaP-TRAP eliminates the confounding effects of transcriptional regulation associated with MPRAs that introduce reporters as DNA. When quantifying the regulatory potential of the 5’ UTR, transcriptional bias may be more pronounced given the region’s proximity to the promoter sequence.

Previous studies have assumed that the frequency of a given Kozak sequence within the genome correlates strongly with its effect on translation.^45^ While this hypothesis has been challenged by MPRAs performed in cell culture, to date we lack measurements of the effect of Kozak strength on translation in zebrafish.^23^ To demonstrate the capacity of NaP-TRAP to quantify the translation of thousands of mRNA reporters simultaneously, we measured the Kozak strength in zebrafish embryos. In support of this approach, we observed a weak correlation between Kozak strength and an *in silico* derived Kozak score (Figure 2E). While our approach has identified conserved activators and repressors (-3 A and -3/-2 U/G) of translation, we have also characterized novel Kozak sequences that differ from the zebrafish or vertebrate consensus Kozaks, including the activating effect of adenosine in the +4 position. Further, we have utilized our random forest regression model to predict the Kozak strength of all zebrafish Kozak sequences (positions -6 to +4). (Table S2). This prediction will improve our annotation of Kozak strength in the zebrafish, as well as our capacity to modulate protein output.

Development requires dynamic spatial and temporal control of translation. Using NaP-TRAP we have identified 5’ UTR *cis*-regulatory elements that differentially regulate translation in development. For example, we show that the zygotically expressed microRNA miR-430 represses the translation of reporters containing miR-430 seeds in the 5’ UTR. Our results are consistent with the *in vitro* observa-tions by *Lytle et al*. in the 5’ UTR,^51^ and support a physiological role for miRNA mediated regulation of 5’ UTRs during developmental transitions *in vivo*.

We have also identified a translation switch driven by C-rich motifs. At 2 hpf C-rich k-mers suppress translation, whereas at 6 hpf these k-mers activate translation. We propose two potential mechanisms to explain this novel 5’ UTR mediated translation control (Figure 6B). First, the expression or loss of a *trans*-acting factor (TAFs) that binds pyrimidine rich tracts may drive differential translation. Members of the poly-pyrimidine tract-binding protein family (PTBP) and components of the EIF3 complex have been shown to activate translation by binding pyrimidine tracts in the 5’ UTR.^52,53^ While future work is necessary to characterize the function of these and other regulators *in vivo*, components of the EIF3 complex have been implicated in the realization of lineage specific translation programs in early embryogenesis.^54^ Second, recruitment of *trans*-factors depends not only on the presence of a given *cis*-regulatory element, but rather the composition of the transcriptome and the pool of available *trans*-factors shapes the regulatory potential of a given element. We observe that the relative importance of U-rich sequences depends on the global rate of translation. When translation is low, the prevalence of U-rich sequences is a prominent predictor of translation (Figure S3D-F). In contrast, as translation increases, the relative importance of U-rich sequences declines, whereas the relative importance of uORFs increases (S3G-I). We propose that this change reflects a change in the availability of the translation initiation machinery.^43^ We speculate that when the supply of ribosomes is limited, the capacity of the 5’ UTR to recruit ribosomes drives translation. In contrast, as the ribosome pool increases, the effect of 5’ UTRs on ribosome recruitment diminishes (Figure 6B).

This competition model can also be employed to explain the differential effect of C-rich tracts on translation. In the early embryo, reporters enriched in U-repeats may recruit the limited supply of ribosomes more efficiently than reporters enriched in C’s. As the supply of translation machinery increases, the effect of competition diminishes, result-ing in the efficient initiation of C-rich 5’ UTRs (Figure 6B). Repressive *trans-*acting factors (TAFs) may amplify the effect of competition, as in the absence of scanning 40S ribosomes, these factors can be more readily recruited to the 5’ UTR. Work from Xiang and Bartel investigating the strong correlation between translation efficiency and poly-A tail length exemplifies this model.^55^ More specifically, they report that prior to gastrulation the pool of PAPBC1 (poly(A) binding protein cytoplasmic 1) is limited, driving competition between transcripts for PAPBC1 occupancy. Transcripts able to bind PAPBC1 have an advantage in recruiting ribosomes from a limited supply of active ribosomes leading to increased translation compared to other transcripts. This competition is eliminated following gastrulation as the relative abundance of PAPBC1 and active ribosomes increases following zygotic genome activation, resulting in the decoupling of poly-A tail length and translation efficiency.

NaP-TRAP also identifies conserved regulators of translation. In both the developing zebrafish embryo and HEK293T cells, poly-U repeats activate translation whereas uORFs repress translation. Interestingly, work from Zinshteyn et. al has demonstrated *in vitro* that yeast EIF4G binds U-repeats greater than five nucleotides, driving translation activation.^56^ While we do not know if EIF4G recruitment modulates translation activation in our experiments, the fact that U repeats are activators of translation across timepoints and experimental systems suggests that this observation may be driven by a component of the canonical translation initiation machinery.

In future works, through the incorporation of different tags in each frame, we propose that NaP-TRAP can be adapted to measure translation in multiple frames simultaneously (Figure 8A-F). We also postulate that NaP-TRAP will enable the field to identify the regulatory potential of both the 5’ and 3’ UTRs across numerous cell types and cellular states. Through this approach, we will not only characterize the regulatory potential of sequences, but also determine how sequence variation in non-coding regions affects human health and disease.

**Figure 8.**
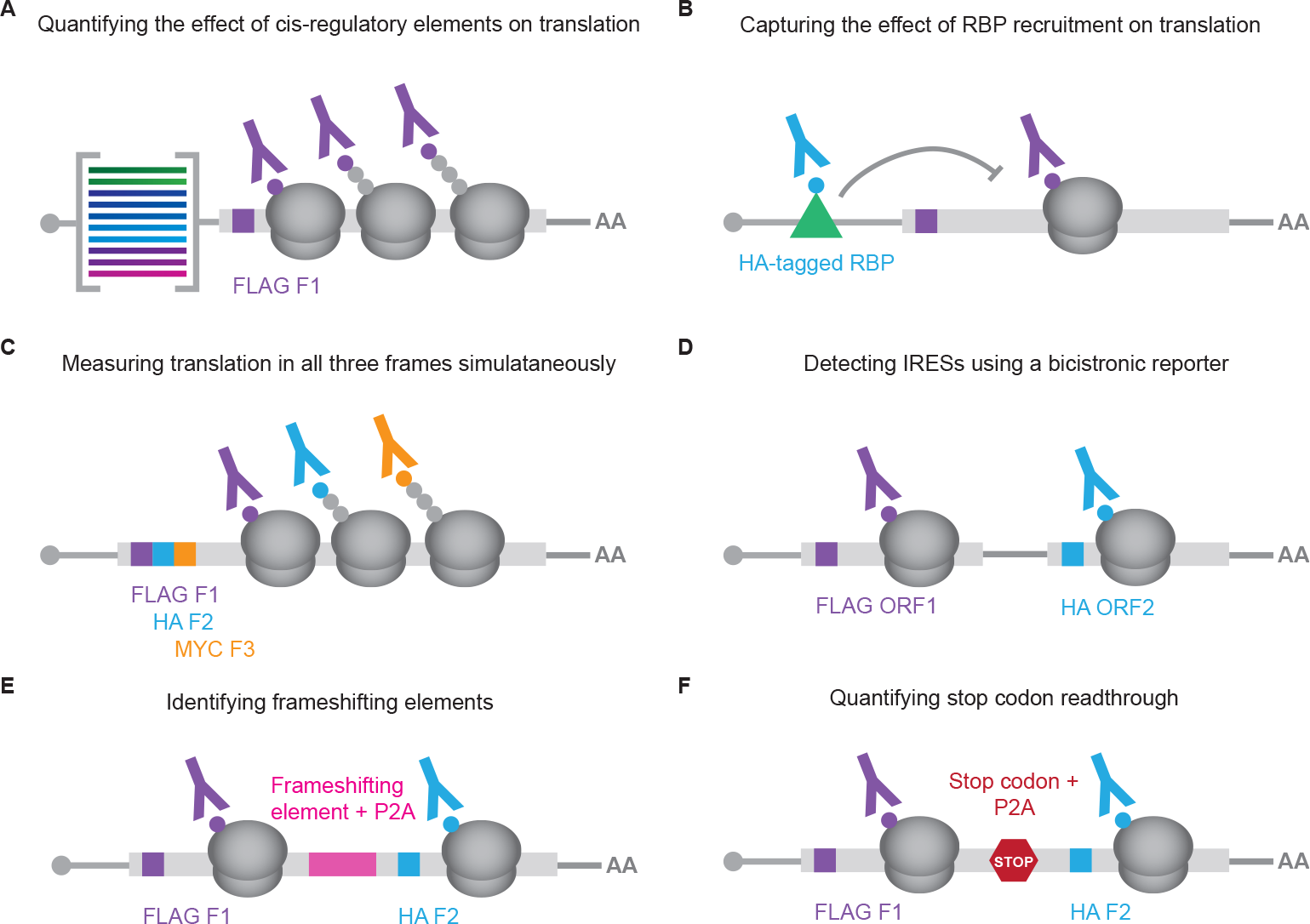
The NaP-TRAP method can be adapted to study the translation of multiple ORFs simultaneously *in vivo*. (A) NaP-TRAP is an accessible, versatile, and quantitative method that measures the translation of thousands of reporters simultaneously through the immunocapture of FLAG-tagged nascent peptides. (B) Through the over-expression of an HA-tagged RBP. NaP-TRAP can be employed in conjunction with an RNA immunoprecipitation experiment to measure the effect of RBP recruitment on translation. (C-F) NaP-TRAP quantifies translation in a frame-specific manner. Through the incorporation of additional epitope tags in frames 2,3 or ORFs outside of the main open reading frame, the NaP-TRAP method can be utilized to: (1) measure out-of-frame translation in the main ORF (C), (2) detect IRES sequences in an unbiased manner through the use of a bicistronic reporter (D), (3) identify frameshifting elements (E), and (4) quantify stop codon readthrough (F).

## Acknowledgements

We thank C. Castaldi, E. Sykes, B. Sullivan, and B. De Kumar from the Yale Center for Genome Analysis for sequencing support; S. Dube, T. Gerson and D. Karpel for technical help; D. Musaev, L. Weiss, S. Hiley, and I. Carmi for feedback on the manuscript; and all members of the Giraldez lab for feedback and support. indent Funding: National Institutes of Health grant R01 HD100035 (A.J.G.), National Institutes of Health grant R35 GM122580 (A.J.G.), National Institutes of Health grant R00 R00HD093873 (J.-D.B.), and National Institutes of Health grant R35 R35GM146883 (J.-D.B).

## Author Contributions

J.-D.B. conceived the NaP-TRAP principle. E.C.S., J.-D.B., and A.J.G. designed the project. E.C.S. and J.-D.B performed the experiments. E.C.S. developed the pipeline to process and analyze NaP-TRAP data. C.E.V. advised E.C.S. on computational methods and supplied reading trimming and demultiplexing software. E.C.S. and A.J.G. wrote the paper. S.K. and H.L. assisted with zebrafish injections and provided experimental support. J.-D.B., S.K. and H.L. provided feed-back on the paper.

## Declaration of Interests

E.C.S., J.-D.B., S.K., and A.J.G are inventors on a provisional patent application (number will be provided at the time of submission) filed by Yale University with the US patent office covering the NaP-TRAP method and the sequences described here.

## Supplemental Information

**Document S1**. Figures S1-S7 (provided in text in the initial submission)

**Supplementary Table 1**. Oligonucleotide sequences related to STAR methods and library Sequences.

**Supplementary Table 2**. Translation data and analysis from NaP-TRAP Kozak Library

**Supplementary Table 3**. Translation data and analysis from NaP-TRAP 5’ UTR Library

**Supplementary Table 4**. Translation data and analysis from NaP-TRAP Validation Library

## Methods

### Resource Availability

#### Lead Contact

Further information and requests for resources and reagents should be directed to and will be fulfilled by the Lead Contact, Antonio J. Giraldez.

#### Materials Availability

Plasmids generated in this study are available from the Lead Contact on request.

### Experimental model and study participant details

#### Zebrafish maintenance and mating

Wild-type zebrafish embryos were obtained through natural mating of TU-AB strain of mixed ages (5-18 months). Mating pairs were randomly chosen from a pool of 60 males and 60 females allocated for each day of the month. Fish lines were maintained following the International Association for Assessment and Accreditation of Laboratory Animal Care research guidelines and approved by the Yale University Institutional Animal Care and Use Committee (IACUC).

#### Hek293T cells

HEK293T cells were grown in a media consisting of Dulbecco’s Modified Eagle Medium (DMEM) (ThermoFisher Scientific #10569010), 10% heat inactivated Fetal Bovine Serum (FBS) (ThermoFisher Scientific #16140071), 20 mM HEPES (ThermoFisher Scientific #15630080), 2 mM L-Glutamine (ThermoFisher Scientific #25030081), 1x Penicillin/Streptomycin (ThermoFisher Scientific #15140122) at 37°C and 5% CO_2_. mRNA transfections were performed using Lipofectamine MessengerMAX Transfection Reagent (ThermoFisher Scientific #LMRNA008) in accordance with the manufacture’s protocol. See Supplementary Methods S1 for a detailed protocol.

## Method details

### NaP-TRAP reporter controls

To enable nascent chain immunocapture, a 3xFLAG tag was incorporated after the first 18 nucleotides of GFP-3xAID* (auxin inducible domain) using an In-Fusion® HD Cloning kit (Takara #638946) (F: 3xFLAG_inf_fwd; R: AID_inf_rev). While AID* domains were included in the initial NaP-TRAP vector to enable future use of an auxin-inducible degron system, this system was not utilized in this study. The vector also included an SP6 promoter sequence and SV40 poly-adenylation signal. NaP-TRAP reporter and dsRED control (plasmid pCS2+-dsRED) mRNAs were generated using a mMESSAGE mMACHINE SP6 transcription kit (Invitrogen #AM1340) from linearized reporter plasmids via NotI-HF® (NEB #R3189L) restriction enzyme digest. *In vitro* transcribed mRNAs were purified using a Monarch® RNA Cleanup Kit (NEB #T2040L) prior to injection.

To validate NaP-TRAP, two reporter experiments were performed. First, to quantify morpholino mediated translation repression using NaP-TRAP, 100 pg of 3xFLAG-GFP-3xAID mRNA and 75 pg of dsRED mRNA were injected into single cell zebrafish embryos in the presence or absence of a morpholino targeting the start codon of 3xFLAG-GFP (250 mM, GFP-MO 5’-AC AGCTCCTCGCCCTTGCTCACCAT-3’, Gene Tools LLC). Embryos were collected at 6 and 24 hpf (25 embryos per NaP-TRAP replicate). Embryos collected for NaP-TRAP were flash frozen in liquid nitrogen prior to sample processing. NaP-TRAP was performed at 6 hpf whereas immunofluorescence was measured at 24 hpf. Images were quantified using ImageJ.^52^ Second, to assess the capacity of NaP-TRAP to measure microRNA mediated repression in the 3’ UTR, two additional reporters were generated: (1) 3xFLAG-GFP-3xmiR-430 and (2) 3xFLAG-GFP-3xmiR-204. Three binding sites of either miR-430 or miR-204 were cloned into the 3’ UTR of 3x-FLAG-GFP-3xAID*, using an In-Fusion® HD Cloning kit (Takara #638946) (F: FGFP_inf_fwd; R: 3xmir430_inf_rev and 3xmir204_inf_rev, respectively). Single cell embryos were injected with 20 pg of both the 3xFLAG-GFP-3xmiR-430 and 3xFLAG-GFP-3xmiR-204 mRNAs, as well as 160 pg of dsRED mRNA. Twenty-five embryos per replicate were collected and flash frozen in liquid nitrogen at 2 and 4.3 hpf.

*For methods detailing NaP-TRAP and qPCR translation measurements see sections: (1) NaP-TRAP (Nascent Peptide Translating Ribosome Affinity Purification) and (2) NaP-TRAP qPCR analysis, respectively.

### NaP-TRAP reporter library assembly

To eliminate excess cytoplasmic 3xFLAG-GFP, a C-terminal PEST domain was incorporated into the NaP-TRAP reporter plasmid using an In-Fusion® HD Cloning Kit (Takara #638946). The 3xFLAG-GFP vector was amplified from the 3xFLAG-GFP-3xAID* plasmid (F: GFP_dd1_inf_fwd, R: GFP_dd1_inf_rev), whereas the PEST domain insert was generated using PCR overlap extension (F: PEST_fwd, R: PEST_rev). For the sake of brevity, 3x-FLAG-GFP-PEST is referred to as 3xFLAG-GFP in the text and figures of this manuscript unless stated otherwise.

NaP-TRAP reporter libraries were constructed using three different PCR reactions:

First, the common coding sequence and 3’ UTR of the reporters were amplified from the 3xFLAG-GFP plasmid, using a forward primer targeting the N-terminus of GFP and a reverse primer targeting the 3’ end of the 3’ UTR (F: 3xFLAG_GFP_fwd, pA-R: pcr_II_pA_rev, sv40-R: sv40_rev). For the Kozak library, a forward primer targeting the sequence immediately downstream of the variable Kozak sequence was used (F: ntrapK_GPF_fwd). All PCR amplicons were gel purified using a Monarch® DNA Gel Extraction Kit (NEB #T1020L).

Second, reporter libraries were amplified from 1 ng of single stranded DNA oligo pools using KAPA HiFi HotStart ReadyMix (Roche #7958935001) for 10-20 cycles of 98°C, 60°C, and 72°C for 15, 20, and 30 seconds, respectively (F: SP6_II_adapt, R: GFP-aug_rev). For initial Kozak library generation see Random Kozak Library section. Each product was PCR purified using a DNA Clean & Concentrator-5 (Zymo #D4014).

Third, to generate a template for *in vitro* transcription, a PCR overlap extension was performed between the reporter library and the purified 3xFLAG-GFP-PEST amplicon using KAPA HiFi Hot-Start ReadyMix (Roche #7958935001) (0.1-1 ng of template DNA). After 10 cycles of 98°C, 60°C, and 72°C for 15, 20, and 30 seconds respectively, primers targeting the 5’ SP6 promoter sequence and the 3’ end of the 3xFLAG-GFP-PEST amplicons were added (F: SP6_II_adapt, pA-R: pCS2_3utr_60A, sv40-R: sv40_rev) followed by an additional 20 cycles at the same conditions. Unless stated explicitly, reporter libraries contained a 60A tail (pA-R: pCS2_3utr_60A). For libraries with an SV40 polyadenylation signal, a reverse primer targeting the 3’ end of the SV40 poly-adenylation signal was used for the amplification of 3xFLAG-GFP-PEST and the assembly of reporter library (R: SV40_rev).

Lastly, templates for *in vitro* transcription were gel purified using a Monarch® DNA Gel Extraction Kit (NEB #T1020L). Reporter mRNAs were generated using a mMESSAGE mMACHINE SP6 transcription kit (Invitrogen #AM1340). *In vitro* transcribed mRNAs were purified using a Monarch® RNA Cleanup Kit (NEB #T2040L). All libraries were injected into single cell zebrafish embryos at 20 pg per embryo.

### Random Kozak library

The random Kozak library consisted of an I5 Illumina adaptor (5’-CCCTACACGACGCTCTTCCGATCT-3’) followed by the 5’ UTR of *Xenopus* beta-globin, seven random nucleotides, six upstream and one downstream of the start codon, and the N-terminus of 3xFLAG GFP. The Kozak library was generated by performing a PCR overlap extension using KAPA HiFi HotStart ReadyMix (Roche #7958935001) for 10 cycles of 98°C, 60°C, and 72°C for 15, 20, and 30 seconds respectively (F: kozak_ntrap_fwd, R: kozak_7_nt_rev, 1 µL of each primer at 100 uM).

### Zebrafish 5’UTR library

A custom single stranded DNA oligo pool consisting of 11,088 oligos was ordered from GenScript (12 K oligo pool). Each 170 nt oligo contained an Illumina I5 (5’-CCCTACACGACGCTCTTCCGA TCT-3’) adaptor sequence, a 124-nucleotide variable region, and 22 nt region with homology to the Kozak sequence and N-terminus of 3xFLAG-GFP (5’-GTAAACATGGTGAGCAAGGGCG-3’). The variable region of the 5’ UTR library was generated using a custom script, tiling the 5’ UTRs of 1,775 maternally supplied genes and six IRES sequences (human AQP4, human MYT2, human NRF, human XIAP, EMCV and crTMV) in 124 nucleotide segments every 25 nucleotides.

### Validation library (Tetramer repeats)

A custom single stranded DNA oligo pool consisting of 256 oligos was ordered from Twist Bioscience as part of a 12 K oligo pool. The design of the common regions of library were identical to that of the zebrafish 5’ UTR library. The variable region (124 nucleotides) consisted of repeats of all possible tetramers. Each repeat occurred 21 times and was separated by a dinucleotide spacer. The dinucleotide spacers were repeated in a pattern across the variable region (TC, AC, AG, CG). These dinucleotide spacers were selected to prevent the creation of unintended upstream ORFs.

### NaP-TRAP spike-ins

The design of the spike-in reporters was identical to that of the zebrafish 5’ UTR and tetramer validation libraries. Each spike-in reporter contained a 20 nucleotide identifier (see below). NaP-TRAP spike-ins were generated by performing a PCR overlap extension using KAPA HiFi HotStart ReadyMix (Roche #7958935001) (F: ntrap_sp1_fwd, ntrap_sp2_fwd, ntrap_sp3_fwd, ntrap_sp4_fwd, ntrap_sp5_fwd; R: ntrap_sp_rev) (1 µL of each primer at 100 µM) for 10 cycles of 98°C, 60°C, and 72°C for 15, 20, and 30 seconds, respectively. Amplicons were gel purified using a Monarch® DNA Gel Extraction Kit (NEB #T1020L). Next, the purified amplicons were cloned into the 3xFLAG-GFP-PEST vector using In-Fusion® HD Cloning (Takara #638946) (vector F: ntrap_spike_inf_fwd, R: ntrap_spike_inf_rev). To gen-erate spike-in mRNAs, plasmids were amplified using KAPA HiFi HotStart ReadyMix (Roche #7958935001) (F: Sp6-Il-adapt, R: pCS2_3utr_60A) for 20 cycles at 98°C, 60°C, and 72°C for 15, 20, and 30 seconds, respectively. Products were then gel purified with a Monarch® DNA Gel Extraction Kit (NEB #T1020L) and then *in vitro* transcribed with a mMESSAGE mMACHINE SP6 transcription kit (Invitrogen #AM1340). mRNAs were purified with a Monarch® RNA Cleanup Kit (NEB #T2040L) prior to use. Spikeins were pooled and added at the RNA extraction step at concentrations of 1, 5, 25, 50, and 125 fg for the 5’ UTR library.

Spike-in #1 TGACGTGGAAGTCGGTCAAG

Spike-in #2 GTCCAGAGACAAAGTCCGGG

Spike-in #3 CACGAGGAGGAACCAGTGAC

Spike-in #4 CTGTTGTTGTGTGAAGGGCG

Spike-in #5 GCTCTCGGTCTCGGAAGAAG

### NaP-TRAP (Nascent Peptide Translating Ribosome Affinity Purification)

To capture tagged nascent chains of reporter mRNAs, an immunoprecipitation using anti-FLAG magnetic beads was performed. Magnetic beads were purchased from three suppliers during the course of the study due to supply chain issues (ANTI-FLAG® M2 Magnetic Beads, Millipore® #M8823; Anti-Flag Magnetic Beads, BioTools LLC #B26102; Pierce Anti-DYKDDDDK Magnetic Agarose, Thermo Fisher Scientific #88836). For the zebrafish and human cell experiments 10 µL and 20 µL of beads (binding capacity of >0.8 mg of FLAG peptide / mL) were utilized, respectively. The magnetic beads were washed with 800 µL of wash buffer three times prior to being added to the lysis solution.

Briefly, frozen embryos and HEK293T cells were lysed in 500 µL of lysis buffer. After 10 minutes at 4°C the lysate was passed through a 25-guage needle (5-10 times). Next, the sample was centrifuged at 16,000 g for 5 minutes at 4°C. The supernatant was transferred to a new Eppendorf tube and 2 µL of DNase I (NEB #M0303L) was added. After a 15 minute incubation at 4°C, the samples were diluted to 1 mL using additional lysis buffer and 75 µL of lysate was collected from each sample to serve as an input. Samples were placed on a rotator at 4°C for 2 hrs. Following incubation and bead capture, the beads were washed with 800 µL of wash buffer three times. The beads (pulldown) and the inputs were then resuspended in 1 mL of Trizol (Invitrogen #15596-018). For the zebrafish 5’ UTR and tetramer validation libraries, spike in reporters were added. RNA extractions were performed in accordance with the manufacture’s protocol. RNA pellets were resuspended in 11 µL of nuclease free H_2_O.

### Library preparation for Next-generation sequencing

Reverse transcription primers (4 µM total) targeting the N-terminus of 3xFLAG GFP were added to purified input and pulldown RNAs (5’-GTGACTGGAGTTCAGACGTGTGCTCTTCCGATCT-CTAC-10N

UMI-TAAC-6 nt sample barcode-GCGTCATGGTCTTTGTAGTCCTC-3’; see NaP-TRAP barcode RT primers in Supplementary Table 1). These primers contained a 6 nucleotide sample barcode and a 10 nucleotide Unique Molecular Identifier (UMI) to allow for demultiplexing and read deduplication, as well as a 3’ I7 Illumina adaptor sequence. To increase library complexity the primer pairs were staggered by 1 nt (e.g., ntrap_RT_b1.1, ntrap_RT_b1.2). Reverse transcription was performed using the Superscript III kit (Invitrogen #18080044) in accordance with manufacturer’s instructions. Reverse transcription reactions were performed at 55°C. cDNA from replicates were pooled and purified by adding AMPure XP Reagent (Beckman Coulter #A63881) at 1.8x the original sample volume. Illumina I5 and I7 forward and reverse primers containing a 10-nucleotide index (see below) were utilized to amplify cDNA libraries via PCR (Kappa Polymerase Master Mix). To reduce the number of PCR duplicates 12-18 cycles were utilized. Amplicons were purified by adding AMPure XP Reagent (Beckman Coulter #A63881) at 0.9x the volume of the PCR reaction. Libraries were sequenced on Illumina NovaSeq 6000 platform.

Illumina I5 primer:

5’-AATGATACGGCGACCACCGAGATCTACAC-10 nt index-ACACTCTTTCC

CTACACGACGCTCTTCCGATCT-3’

Illumina I7 primer:

5’-CAAGCAGAAGACGGCATACGAGAT-10 nt index-GTGACTGGAGTTCAGA

CGTGTGCTCTTCCGATCT-3’

### NaP-TRAP-qPCR analysis

Following NaP-TRAP, cDNA was synthesized using random hexamers (ThermoFisher Scientific #N8080127) and SuperScript III Reverse Transcriptase (Invitrogen #18080044) following the manufacturer’s protocol. Translation values were determined by qPCR using the Power SYBR Green PCR Master Mix (Applied Biosystems #4367659). Levels of reporters in the input and pull-down were normalized using the 2-ΔΔCt method,^53^ where the NaP-TRAP reporter was the target and dsRED the control. Translation was calculated as a ratio of fold enrichment in the pulldown relative to the input.

Primers utilized in qPCR:

1. Translation blocking morpholino (GFP_qpcr_fwd, GFP_qpcr_rev)
2. MicroRNA seeds in 3’ UTR (3xmir-430_qpcr_fwd, 3xmir-204_qpcr_fwd, T7_qpcr_rev)
3. U/C rich reporters (igf1ra_wt_qPCR_fwd, igf1ra_mut_ qPCR_fwd, asun_wt_qPCR_fwd, asun_mut_qPCR_fwd, sich211_wt_qPCR_fwd, sich211_mut_qPCR_fwd, atp6v0_ wt_qPCR_fwd, atp6v0_mut_qPCR_fwd, GFP_qPCR_5p_rev)
4. dsRED (dsRED_qpcr_fwd, dsRED_qpcr_rev)

### NaP-TRAP NanoLuc reporter construction

We substituted NanoLuc luciferase (amplified from a gBlocks Gene Fragment, Integrated DNA Technologies IDT) for GFP in the 3xFLAG-GFP-PEST vector using In-Fusion® HD Cloning (Takara #638946) (3xFLAG-PEST vector F: PEST_fwd, R: FLAG_rev_inf; NanoLuc insert F: FLAG-Nluc_inf_fwd, R:

Nluc_inf_rev). Note: The PEST domain was included in NanoLuc constructs to reduce the half-life of the NanoLuc protein and thereby improve the sensitivity of the assay. To generate NanoLuc reporters 3xFLAG-NanoLuc-PEST was amplified using KAPA HiFi HotStart ReadyMix (Roche #7958935001) (F: 3xFLAG_GFP_fwd R: pcr_II_pa_rev) and gel purified using a Monarch® DNA Gel Extraction Kit (NEB #T1020L).

### Kozak NanoLuc reporters

To construct Kozak NanoLuc reporters, 3xFLAG-NanoLuc (amplicon from previous section) was amplified with four different forward primers (ntrap_k1_fwd, ntrap_k2_fwd, ntrap_k3_fwd, and ntrap_k4_fwd) and a common reverse primer (pcr_II_pa_rev) using KAPA HiFi HotStart ReadyMix (Roche #7958935001) at 20 cycles of 98°C, 60°C, and 72°C for 15, 20, and 45 seconds, respectively. Products were gel purified using a Monarch® DNA Gel Extraction Kit (NEB #T1020L). To add an SP6 promoter sequence and a 60A hard-encoded tail the purified product (1 ng) was amplified using KAPA HiFi HotStart ReadyMix (Roche #7958935001) at 20 cycles of 98°C, 60°C, and 72°C for 15, 20, and 45 seconds, respectively (F: Sp6-Il-adapt, R: pCS2_3utr_60A). Products were gel purified with a Monarch® DNA Gel Extraction Kit (NEB #T1020L) and then *in vitro* transcribed with a mMESSAGE mMACHINE SP6 transcription kit (Invitrogen #AM1340). mRNAs were purified with a Monarch® RNA Cleanup Kit (NEB #T2040L) prior to injection. Note, the input of one of the replicates of the Kozak reporter K4 was below the read filter cutoff in the NaP-TRAP Kozak Library experiment.

### 5’ UTR miR-430 NanoLuc reporters

4xmiR-430-nanoluc and 4xmiR-430-MUT-nanoluc reporters were generated through PCR overlap extension (WT-F: 4xmiR-430-fwd, WT-R: 4xmiR-430-rev; MUT-F: 4xmiR-430-MUT-fwd, MUT-R: 4xmiR-430-MUT-rev) using KAPA HiFi HotStart ReadyMix (Roche #7958935001) at 10 cycles of 98°C, 60°C, and 72°C for 15, 20, and 30 seconds, respectively (1 µL of each primer at 100 µM). The extension products were then gel purified using a Monarch® DNA Gel Extraction Kit (NEB #T1020L).

Full-length reporters were constructed by performing a PCR overlap extension of the 5’ UTR product with the NanoLuc amplicon described in the Cloning NanoLuc section using KAPA HiFi HotStart ReadyMix (Roche #7958935001). After 10 cycles of 98°C, 60°C, and 72°C for 15, 20, and 40 seconds, respectively, primers targeting the SP6 promoter sequence (F: Sp6-Il-adapt), and the 3’ end of the nanoLuc amplicon were added (R: pCS2_3utr_60A). The PCR reaction was continued for an additional 20 cycles at the same cycle conditions. Products were gel purified with a Monarch® DNA Gel Extraction Kit (NEB #T1020L) and then *in vitro* transcribed with ta mMESSAGE mMACHINE SP6 transcription kit (Invitrogen #AM1340). mRNAs were purified with a Monarch® RNA Cleanup Kit (NEB #T2040L) prior to injection.

### Dual luciferase assay

Firefly luciferase was amplified from a pCS2+-Fluc plasmid using KAPA HiFi HotStart ReadyMix (Roche #7958935001) at 20 cycles of 98°C, 60°C, and 72°C for 15, 20, and 50 seconds (F: SP6_ext_fwd, R: SV40_60A_rev). Products were gel purified with a Monarch® DNA Gel Extraction Kit (NEB #T1020L) and then *in vitro* transcribed with a mMESSAGE mMACHINE SP6 transcription kit (Invitrogen #AM1340). mRNAs were purified with a Monarch® RNA Cleanup Kit (NEB #T2040L) prior to injection.

Single cell zebrafish embryos were co-injected with mRNAs encoding nano and firefly luciferase (0.5 pg of nano / 19.5 pg of firefly). Embryos were collected at 6 hpf and frozen in liquid nitrogen (5 embryos per replicate). Nano and firefly luciferase activities were measured using the Nano-Glo® Dual-Luciferase® Reporter Assay System (Promega #N1610). Firefly luciferase activity was utilized to normalize nano luciferase measurement across reporters.

### C and U rich reporters

C- and U-rich wild-type and mutant reporters were generated through PCR overlap extension (Primers used: igf1ra_wt_fwd, igf1ra_wt_rev; igf1ra_mut_fwd, igf1ra_mut_rev; asun_wt_fwd, asun_wt_rev; asun_mut_fwd, asun_mut_rev; sich211_wt_fwd, sich211_wt_rev; sich211_mut_fwd, sich211_mut_R atp6v0_wt_fwd, atp6v0_wt_rev; atp6v0_mut_F atp6v0_mut_R) using KAPA HiFi HotStart ReadyMix (Roche #7958935001) at 10 cycles of 98°C, 60°C, and 72°C for 15, 20, and 30 seconds, respectively (1 µL of each primer at 100 µM). The extension products were then gel purified using a using Monarch® DNA Gel Extraction Kit (NEB #T1020L).

Extension products were cloned into the 3xFLAG-GFP-3xAID* vector using In-Fusion® HD Cloning (Takara #638946) (vector F: 3xFLAG_GFP_F R: I5_inf_rev). Plasmids were digested with NotI-HF® (NEB #R3189L) and *in vitro* transcribed with a mMES-SAGE mMACHINE SP6 transcription kit (Invitrogen #AM1340). mRNAs were purified with a Monarch® RNA Cleanup Kit (NEB #T2040L) prior to injection.

## Quantification and Statistical Analysis

### Read trimming and translation calculation

Reporter libraries were sequenced on an Illumina NovaSeq 6000 platform. Sequencing data were stored using LabxDB.^54^ Paired-end reads were trimmed and demultiplexed using Read-Knead (https://github.com/vejnar/ReadKnead). Barcodes identifying replicates and UMIs were extracted from read two. Common regions were trimmed from read one to facilitate accurate mapping. For the Kozak library, reporters were counted using a custom python script. Reporters with indels in the Kozak sequence or reporters without an AUG in the appropriate position were eliminated. For the zebrafish 5’ UTR and validation libraries, reads were mapped to a library specific index using Bowtie2.^55^ PCR duplicates were eliminated using UMIs. UMIs were considered identical if they had a Hamming Distance less than 2. Reads for each experiment were normalized by dividing the read counts of each reporter by the sum of the total number of reads mapped to the spike-ins. In the 5’ UTR reporter library (pA and sv40 in zebrafish) spike-in #3 was eliminated from the analysis, because in some of the samples the read counts for spike-in #3 did not correlate with amount of spike-in added. In the absence of spike-ins, read counts were normalized based on the total number of mapped reads per replicate (reads per million, RPM). Translation values were calculated as a ratio of reads in the pulldown relative to reads in the input. Translation values were only included in the downstream analyses if the input contained greater than or equal to 100 unique reads across all replicates.

### Random Forest Regression models

Random forest regression models were employed to predict translation (scikit-learn 1.3; RandomForestRegressor).^56^ For the Kozak library, features were generated by one-hot encoding positions -6 to -1 and position +4 of each reporter sequence. In contrast, for the 5’ UTR library, k-mer counts (1-6 nucleotides) and uORF features were generated using a custom python script. For the 5’ UTR library, features were filtered by calculating the Spearman Rank Correlation Coefficient (SPR) between each feature and translation prior to model training. Features that had a correlation greater than 0.05 or less than -0.05 were included in the random forest model.

To prevent model overfitting, the data were divided randomly into two groups: a test and training set, comprised of 30% and 70% of the reporters, respectively. To optimize model parameters, the training data were divided into five different groups of equal size and a 5-fold cross-validation was performed. The following parameters were optimized using an exhaustive grid search (n_estimators: 20, 100, 200; max_features: 10, 20 and 30 percent of supplied features; max_depth: 3, 5, 7 and min_samples_split: 2, 4, 8). Bootstrapping was employed to select samples used to train each tree. The predictive power of the model was assessed using the test set. A permuted feature importance analysis was performed to identify the features with the greatest predictive power.

### Differential motif enrichment analysis

Reporters were ranked based on their translation at 2 and 6 hpf. Using the sum and difference of these rankings across timepoints, four groups of reporters were generated: (1) repressed, (2) active, (3) repressed post ZGA (active in HEK293T cells), and (4) active post ZGA (active in zebrafish). Repressed and active reporters constituted the top and bottom 10% of reporters based on the sum of their ranks at 2 and 6 hpf, respectively, whereas the repress post ZGA and active post-ZGA. were the top and bottom 10% of groups based on the difference between their ranks at 2 and 6 hpf. A differential motif enrichment analysis was performed on each group. Fold enrichment values were determined by dividing the count of each k-mer in the reporter group by the count of the k-mer in the library, whereas the significance of the fold-change was determined using a hypergeometric test (Bonferroni corrected p-value threshold).

### miRNA complementarity analysis

miR-430 (GCACUU) and miR-1 (ACAUUC) seeds were identified in the reporter library. The Vienna RNAcofold program^57^ was utilized to measure the complementarity between the section of the reporter mRNA (20 nt upstream and 7 nt downstream of the 5’ end of seed site) and the microRNA. For miR-430 the miRNA species with the highest complementary was selected for downstream analysis.

miR-430a: 5’-UAAGUGCUAUUUGUUGGGGUAG-3’ miR-430b : 5’-AAAGUGCUAUCAAGUUGGGGUAG-3’ miR-430c : 5’-UAAGUGCUUCUCUUUGGGGUAG-3’

miR-1-1 / mir-1-2: 5’-UGGAAUGUAAAGAAGUAUGUAU-3’

### Feature rank analysis

To generate the feature rank plot (Figure 6A), features (k-mer counts of four nucleotides or fewer) were ranked based on the mean difference in translation between reporters enriched and depleted in the feature, top and bottom 20% respectively, at 2 and 6 hpf. Given the prominent effect of upstream open reading frames on translation, reporters with uORFs were excluded from the analysis. Features were only included in the analysis if there was a significant mean difference in translation at either timepoint (Mann-Whitney U test with Bonferroni corrected p-value threshold).

### Statistical analyses

The plots and statistical analyses in Figures 1B, 1C, 2G, 5E and 6C-F were generated using GraphPad Prism. All other analyses unless otherwise stated were performed using custom scripts written in Python 3. Plots were generated using the Matplotlib package.^58^ Venn diagrams were generated using Matplot-venn (https://github.com/konstantint/matplotlib-venn). Statistical analyses were performed using the SciPy^59^ and NumPy^60^ packages, whereas the random forest analysis was performed using the scikit-learn package.^56^ Feature and experimental data were stored using an SQLite database (https://www.sqlite.org/).

## Data and code availability

Raw reads will be made publicly accessible in the Sequence Read Archive at the time of publication (submission currently in progress). All scripts used to process and analyze NaP-TRAP data will be released at the time of publication (https://github.com/ecstrayer/nap-trap_paper).

**Figure S1.**
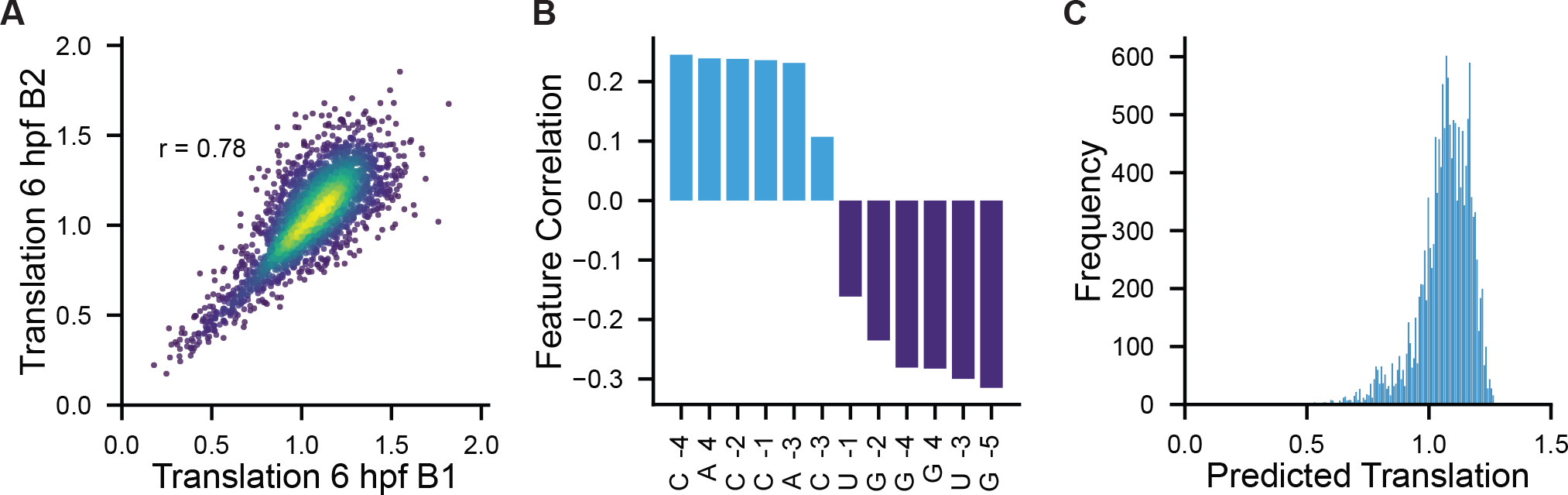
Using NaP-TRAP to model and predict Kozak Strength in early embryos. (A) Plot comparing replicate translation values for the Kozak library (Pearson’s R; N = 2,712 reporters). (B) The correlation coefficient between the top 12 features identified by the permuted feature importance analysis and translation (blue refers to positive correlation, purple refers to negative correlation; Spearman Rank Correlation Coefficient). (C) The predicted translation values of all possible Kozak sequences in early zebrafish embryos (N = 15,364 Kozak sequences) (Random forest regression model: Figure 2B-D) (Table S2).

**Figure S2.**
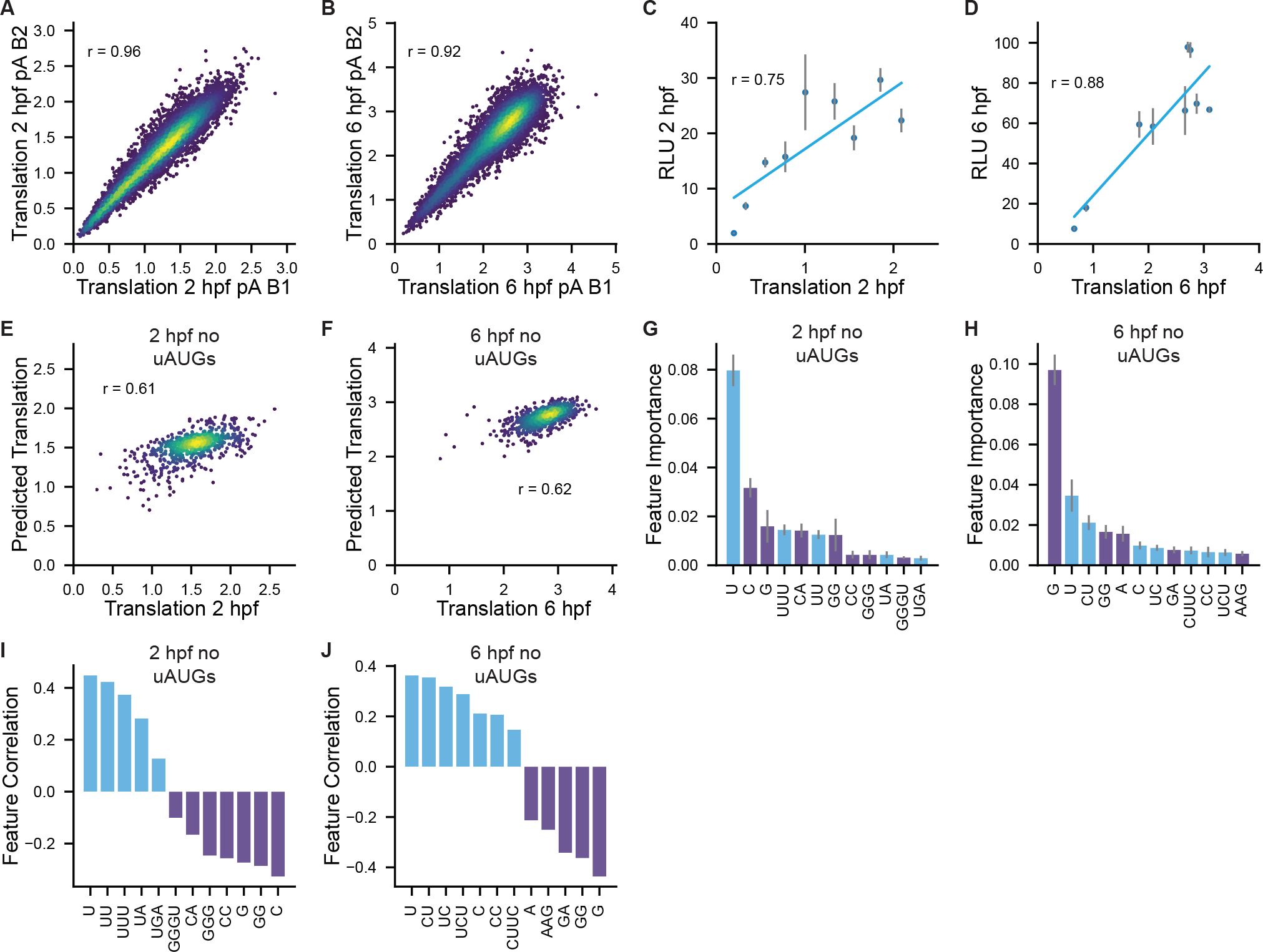
NaP-TRAP quantitatively measures the translation of zebrafish 5’ UTRs. (A and B) Comparing the translation values of replicates of the 5’ UTR library with a hard-encoded poly-A at 2 and 6 hpf (Pearson’s R) (A: N = 9,516 reporters; B: N = 8,529 reporters). (C and D) Random forest regression models were trained on reporters without upstream AUGs. The performance of each model was evaluated by predicting the translation values of a test set of reporters (Pearson’s R) (N = 660 reporters). (E-H) The permuted feature importance of the top 12 features informing the predictive power of each model (E, F). The spearman rank correlations between the top 12 features and translation at each timepoint (G, H) (blue represents positive correlation, purple represents negative correlation).

**Figure S3.**
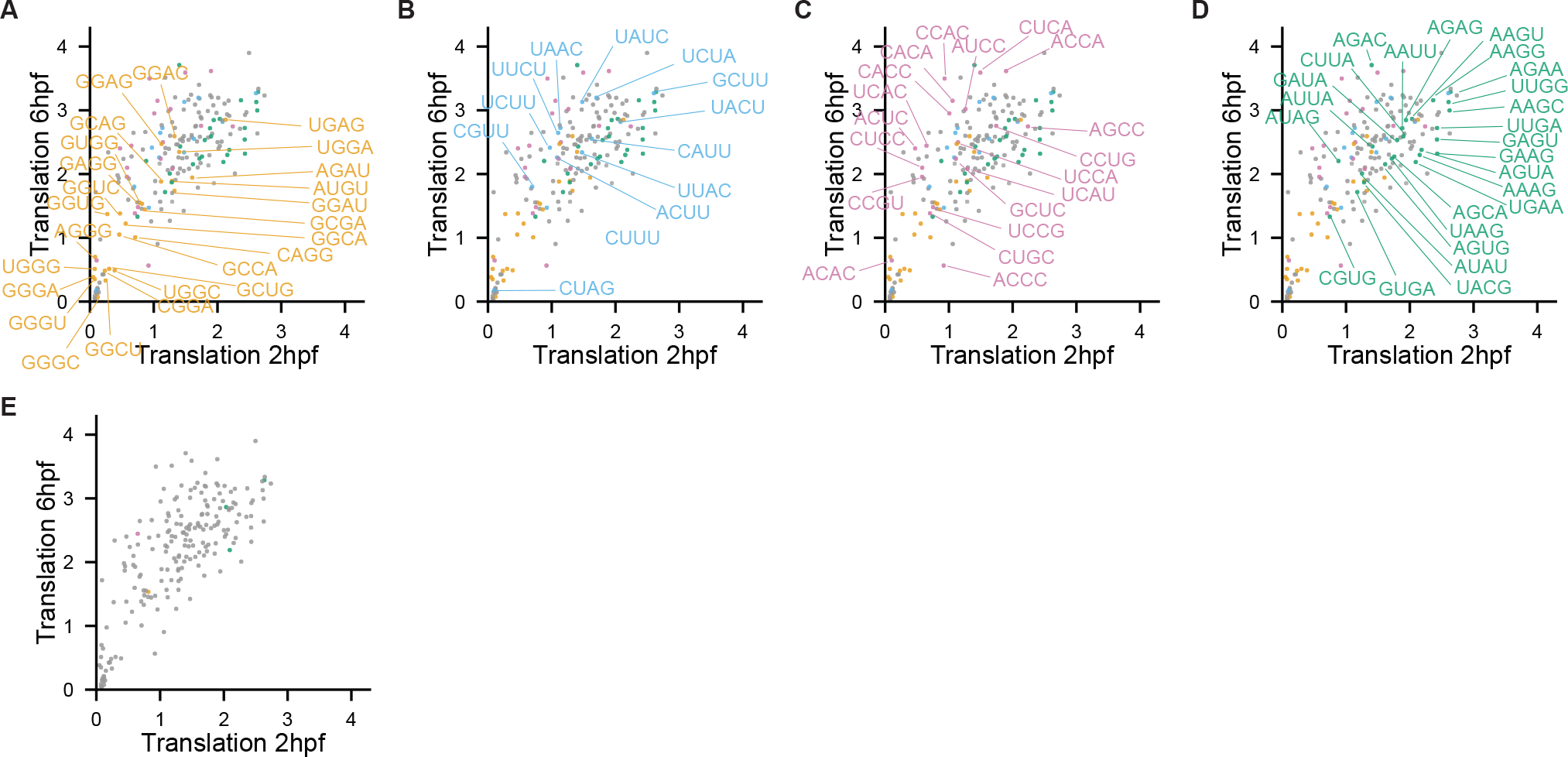
Using NaP-TRAP to measure the regulatory activity of cis-elements in an environment with reduced translation. (A-B) Replicates of translation values for SV40 zebrafish 5’ UTR library (Pearson’s R) (A: N = 9,999 reporters; B: N = 7,442 reporters). (C)The distributions of the translation values for the hard encoded pA (60A) and the sv40 zebrafish 5’ UTR libraries at 2 and 6 hpf (Mann-Whitney U test p < 10^-100^) at 2 and 6 hpf (N = 7,427 reporters). (D-I) Random forest regression models predict translation at 2 and 6 hpf (N = 2,233 reporters) (D, G). We utilized a permuted feature importance analysis to identify the top 12 features informing the prediction of the RFM at both timepoints (E, H). We plotted the Spearman Rank Correlation of each of the selected features with translation (blue is positive and purple is negative) (F, I). (J) NaP-TRAP derived translation values the SV40 zebrafish 5’ UTR library at 2 hpf and 6 hpf. Reporters colored by enrichment group (yellow: repressive, blue: active, pink: active post-ZGA, and green: active pre-ZGA). (K-N) The enrichment for all pentamers in each group relative to all reporters (hyper-geometric test to calculate significance with Bonferroni corrected p-value threshold).

**Figure S4.**
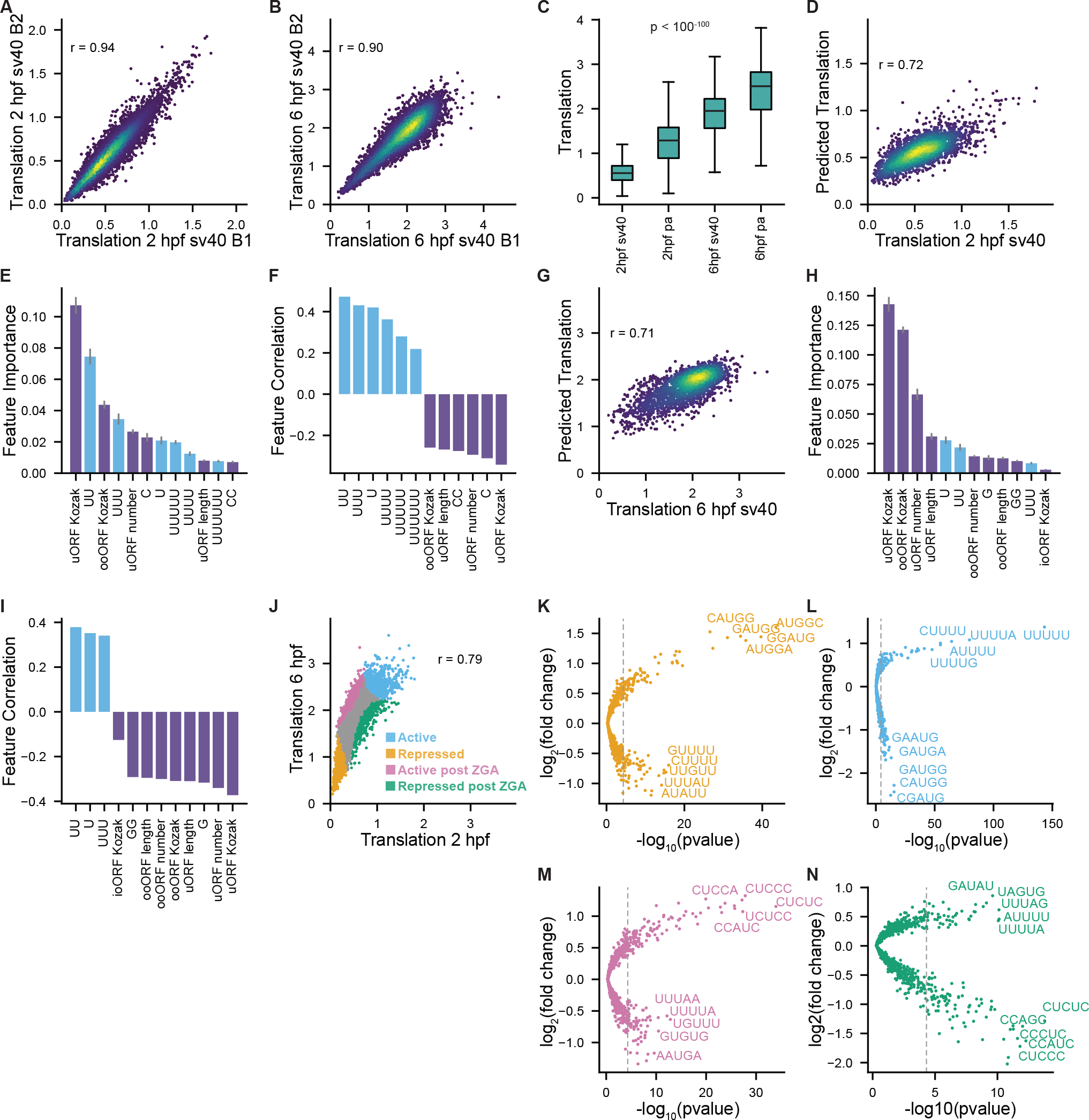
NaP-TRAP validation library of tetramer repeats. (A-D) Reporters were labeled based on whether their encoded tetramer repeat was enriched in the reporter groups defined in the differential kmer enrichment analysis performed on the 5’ UTR library (Figure 4B-E, Table S3) (orange: repressive, blue: active, pink: active post ZGA, green: repressed post ZGA, gray: no group). The validation of the active motifs was limited by the length of the motifs as the dinucleotide spacers created the motifs present in the other groups (B). (E) Previously labelled reporters were colored gray if they contained tetramers found in other groups (four or more counts).

**Figure S5.**
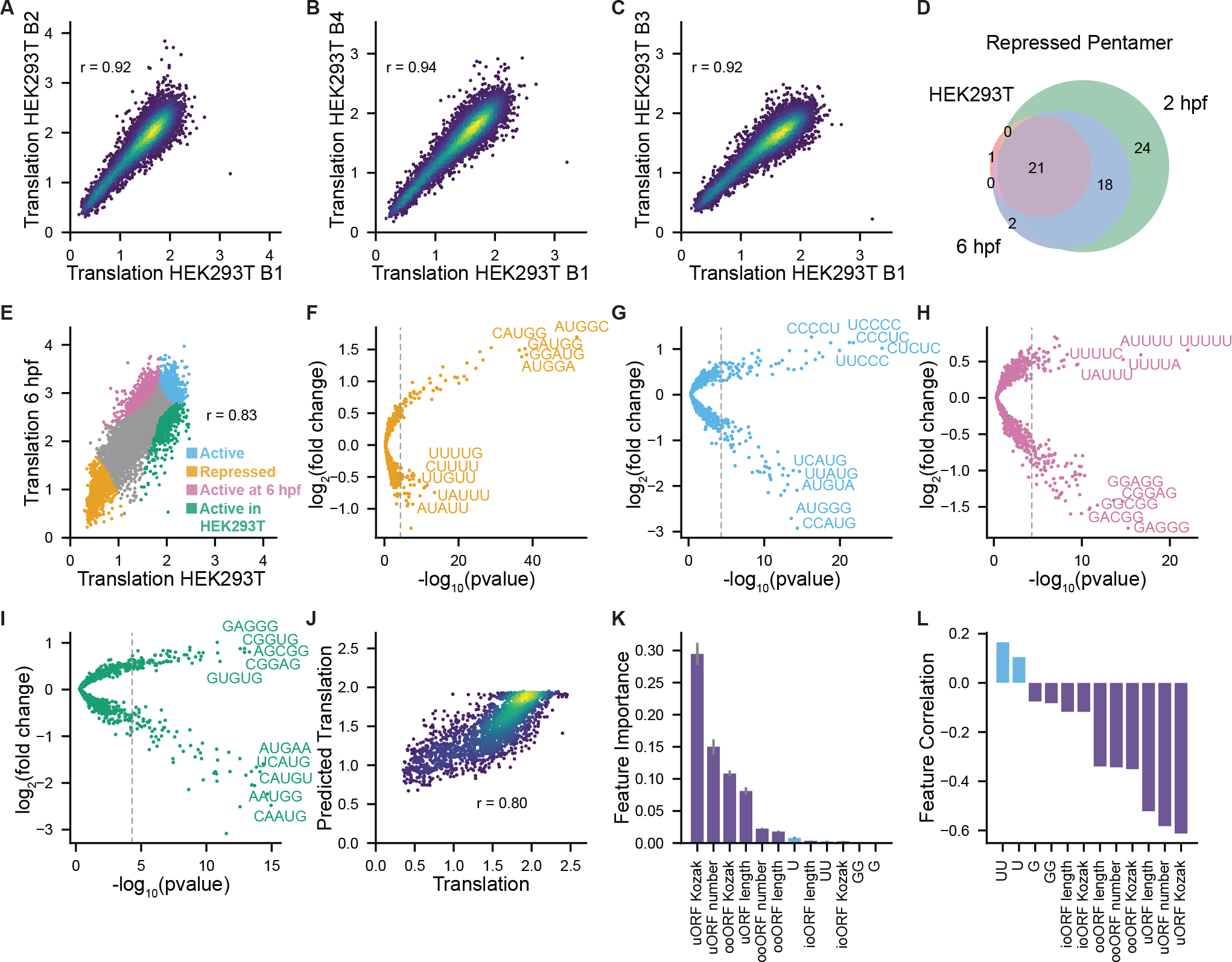
Comparing translation in HEK293T cells and the developing zebrafish embryo. (A-C) Replicates of translation values for the 5’ UTR library in HEK293T cells (Pearson’s R) (N = 7,584 reporters). (D) Venn-diagram comparing the repressive features of HEK293T cells and the zebrafish embryo at 2 and 6 hpf (pentamers enriched in the bottom 10% of reporters). (E) NaP-TRAP derived translation values from the zebrafish embryos at 6 hpf and HEK293T cells at 12 hpt. Both libraries had a hard-encoded 60A tail. Reporters colored by enrichment group (orange: repressive, blue: active, pink: active at 6 hpf, and green: active in HEK293T cells) (N = 7,506 reporters). (F-I) The enrichment for all pentamers in each group relative to all pentamers found in the reporter library. Significance was determined using a hyper-geometric test (Bonferroni corrected p-value threshold). (J-L) Random forest regression model predicts translation (N = 2,252 reporters) (J). The permuted feature importance for features with the greatest effect on the model’s predictive power (top 12; K). The correlation of said features with translation (L) (blue is positive correlation, purple is negative correlation, Spearman Rank Correlation Coefficient)

